# Single-cell mRNA-regulation analysis reveals cell type-specific mechanisms of type 2 diabetes

**DOI:** 10.1101/2023.03.23.533985

**Authors:** J.A. Martínez-López, A. Lindqvist, A. Lopez-Pascual, P. Chen, L. Shcherbina, S. Chriett, N.G. Skene, R.B. Prasad, M. Lancien, P.F. Johnson, P. Eliasson, C. Louvet, A.B. Muñoz-Manchado, R. Sandberg, J. Hjerling-Leffler, N. Wierup

**Author notes:** Co-last author (J. Hjerling-Leffler, N. Wierup).

## Abstract

Perturbed secretion of insulin and other pancreatic islet hormones is the main cause of type 2 diabetes (T2D). The islets harbor five cell types that are potentially altered differently by T2D. Whole-islet transcriptomics and single-cell RNA-sequencing (scRNAseq) studies have revealed differentially expressed genes without reaching consensus. Here, we demonstrate that unprecedented insights into disease mechanisms can be obtained by network-based analysis of scRNAseq data. We developed differential gene coordination network analysis (dGCNA) and analyzed islet scRNAseq data from 16 T2D and 16 non-T2D individuals. dGCNA revealed T2D-induced cell type-specific networks of dysregulated genes with remarkable ontological specificity, thus allowing for a comprehensive and unbiased functional classification of genes involved in T2D. In beta cells eleven networks of genes were detected, revealing that mitochondrial electron transport chain, glycolysis, cytoskeleton organization, cell proliferation, unfolded protein response and three networks of beta cell transcription factors are perturbed, whereas exocytosis, lysosomal regulation and insulin translation programs are instead enhanced in T2D. Furthermore, we validated the ability of dGCNA to reveal disease mechanisms and predict the functional context of genes by showing that *TMEM176A/B* regulates the beta cell cytoskeleton and that *CEPBG* is a key regulator of the unfolded protein response. In addition, comparing beta- and alpha and cells, we found substantial differences, reproduced across independent datasets, confirming cell type-specific alterations in T2D. We conclude that analysis of networks of differentially coordinated genes provides outstanding insight into cell type-specific gene function and T2D pathophysiology.

## INTRODUCTION

Type 2 diabetes (T2D) is a major health care challenge caused by a combination of insulin resistance in target tissues and insufficient insulin release from pancreatic islet beta cells. Without doubt, impaired islet function is central in T2D but the mechanisms behind are not fully understood^1,2^. Transcriptomic data from whole islet preparations have identified differentially expressed genes in T2D^3-7^. A confounding factor in these studies is that islets are composed of five functionally distinct endocrine cell types^8^. Single-cell RNA-sequencing (scRNAseq) studies of islet cells have revealed cell type-specific gene expression alterations, but with little overlap between studies^9-13^. Thus, novel computational analysis and functional validation have been identified as crucial for advancing our understanding of T2D islet disease mechanisms^13,14^. Network biology allows for the identification of pathways and regulatory mechanisms by inferring network modules^15-18^. Disease-related changes can be identified by comparing network structures^19,20^. Differential Network Analysis (DiNA) instead relies on constructing and analyzing the network structure of the dynamics between states^21-24^. Here we developed a version of DiNA specific for scRNAseq data, called differential Gene Coordination Network Analysis (dGCNA). We show that analyzing networks of differentially coordinated genes is a viable and unbiased approach to infer cell type-specific dynamics of the disease from static data. In beta cells, we identified non-canonical genes and pathways in addition to multiple genes and pathways previously implicated in T2D beta cell dysfunction. Importantly, dGCNA also predicted the functional context of genes. Additionally, we identified genes and pathways involved in alpha cells confirming that T2D entails cell type-specific gene regulatory changes.

## RESULTS

In this study we generated and merged two scRNAseq data sets from human pancreatic islets. Data set 1 was generated from hand-picked islets (with confirmed ability to release insulin) from six non-T2D (HbA1c <6.0%) and six T2D (HbA1c>6.5% or diagnosis) brain dead organ donors. Cells were dissociated and analyzed acutely (within 48 hours of collection) using Smart-seq2^25^. After filtering, we analyzed 3645 cells in data set 1. In data set 2 we analyzed 4866 islet cells from 10 non-T2D donors and 10 patients diagnosed with T2D from commercial sources (Prodo Laboratories Inc.) using the same Smart-seq2 pipeline. Part of data set 2 has been published^9^. Characteristics of the 32 donors are presented in Table S1). After merging the data sets, we detected on average 6000 genes in alpha- and beta cells (Fig. S1B). Conos integration and clustering revealed eleven distinct cell types expressing known markers for islet cell types (Fig. 1A-D and Fig. S1A).

**Figure 1.**
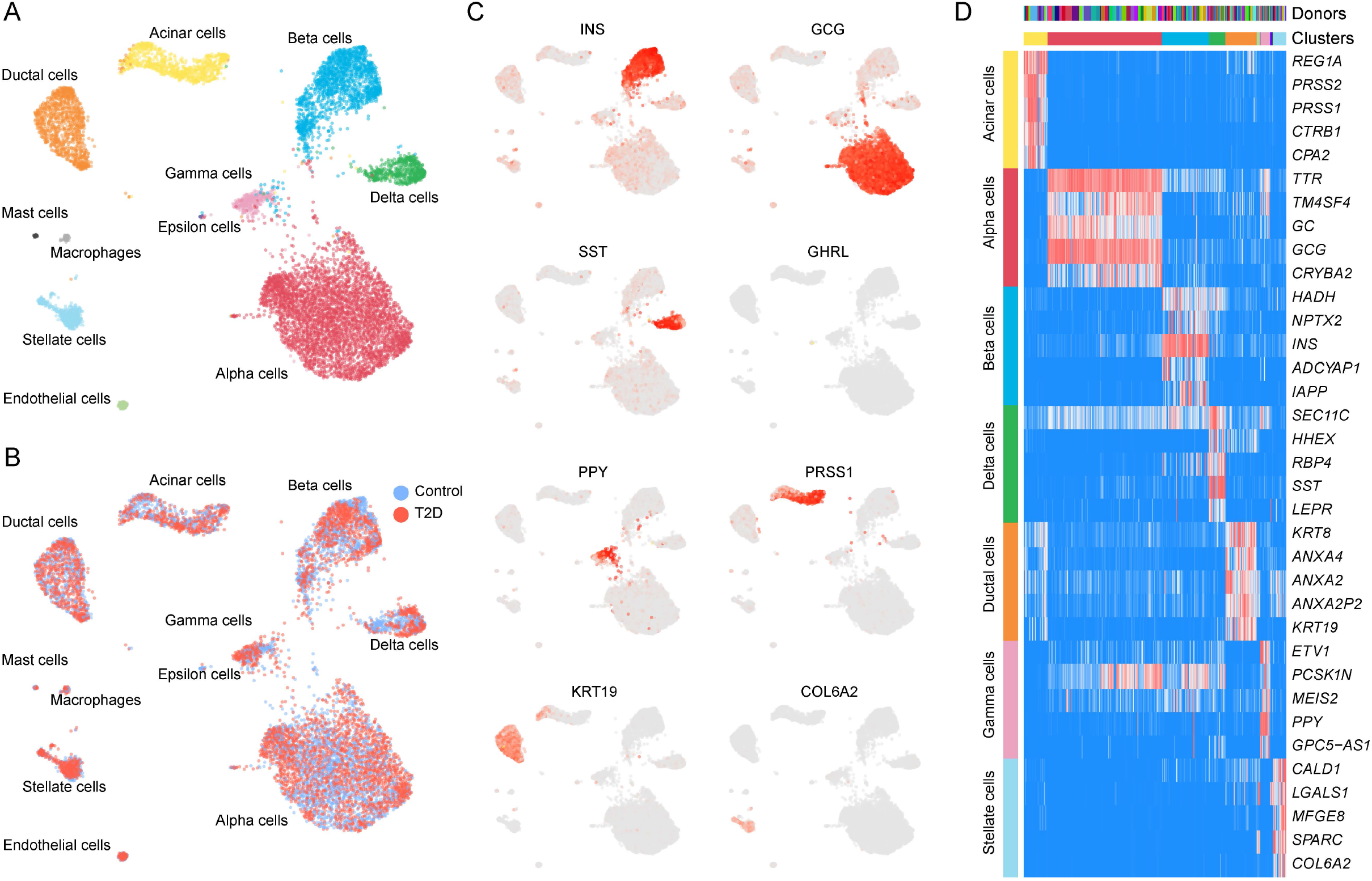
Cell type clustering. UMAP visualization of 8511 single cells, from 32 donors with cell types **(A)** and diagnosis of donors **(B)** indicated. Cell type marker genes indicated in **(C and D)**. See also Fig. S1.

Clustering of beta cells alone (non-T2D and T2D separately) of data set 1 revealed similarities in gene expression patterns (correlations between genes) in non-T2D cells that were lost in T2D cells (Fig. 2A). The identification of such groups of genes suggests that single-cell transcriptomes might represent snapshots of parallel cellular states captured across different phases. Hence, RNA expression could be coordinated in a pathway-specific manner in T2D beta cells. We decided to investigate if these patterns of changes on a whole transcriptome scale could reveal cell type-specific T2D disease biology. To identify the structure of differentially coordinated gene programs that had either been strengthened (hyper-coordinated) or weakened (de-coordinated) in T2D cells, we developed dGCNA, which harnesses the high-dimensionality and biological variability of single-cell transcriptomics (Fig. 2B). In essence this is a differential network analysis on single cell types meaning that the differential network is built on the difference in gene-pair coordinations between the two states (non-T2D and T2D). We performed statistical comparisons of correlation coefficients between gene pairs in a single cell type at a time, using a linear mixed-effect model to account for donor specific effects (Fig. 2B2) and then used a dynamic boot-strap-based threshold to identify robust links to create a robust differential network (RDN) (Fig. 2B3). To analyze the network, we then performed topological analysis and gene clustering on this network (Fig 2B4-B6)

**Figure 2.**
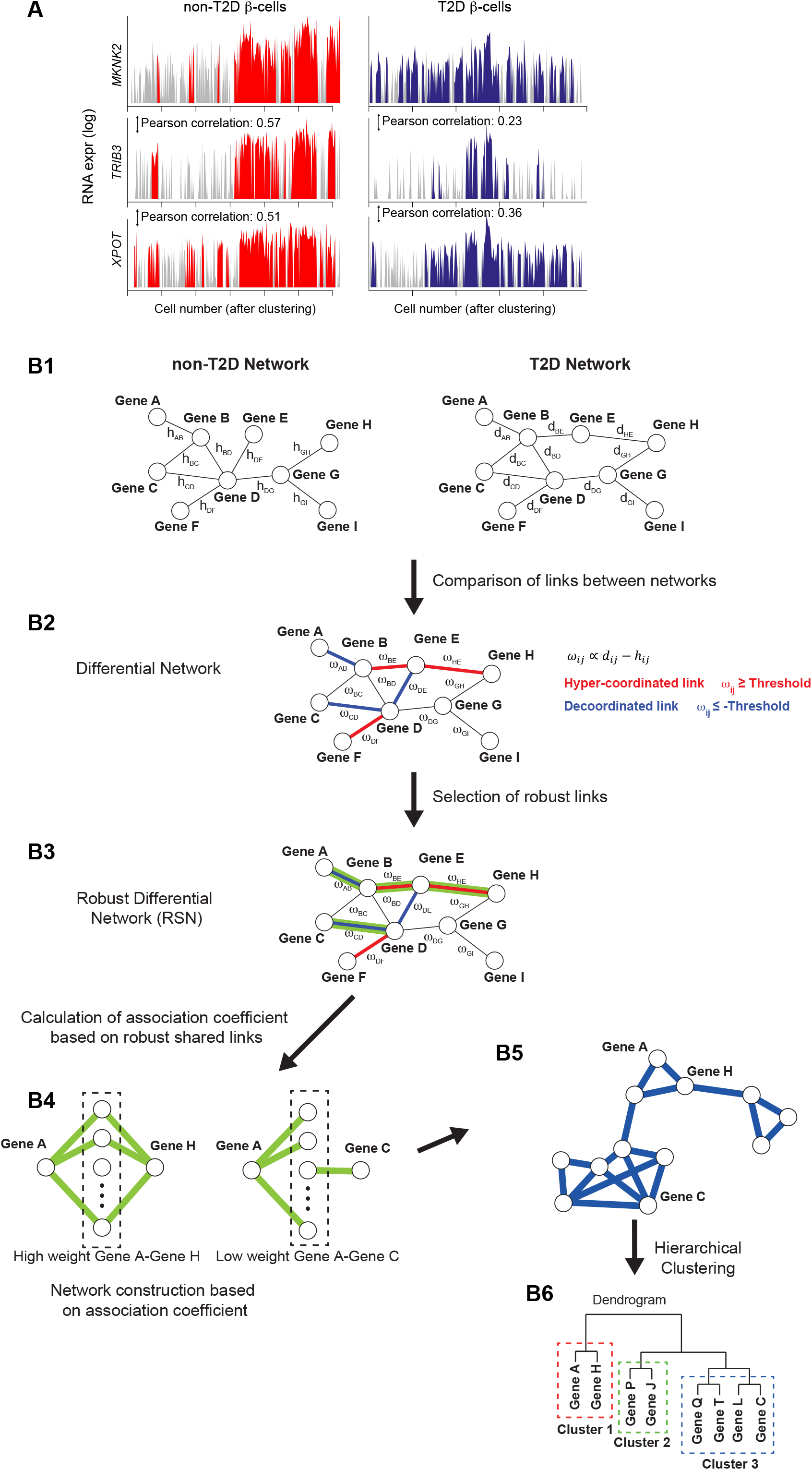
Differential gene coordination analysis. Example of observed changes in gene-gene correlation between non-T2D and T2D donors **(A). (B)** Differential gene coordination approach to reveal disease-related networks in a single cell type. B1: creation of two matrices (one for each state) containing all gene-gene correlations determined as the residuals of a mixed model controlling for donor ID. B2: comparison of the two matrices creating a delta-matrix based on change in correlation. B3: creation of a robust coordination network by selection of robust changes defined as significant based on bootstrapping analysis on random connections. B4: calculation of association coefficients based on robust differential network (RDN) B5: Topological analysis and network construction. B6: Hierarchical clustering of networks.

To reveal T2D-affected biological pathways in beta cells, we performed dGCNA, and the resulting dendrogram indicated a high level of modularity (Fig. 3A), revealing eleven networks of differentially coordinated genes (NDCGs) pointing to pathway alterations in T2D beta cells (Fig. 3A and Table S2). Each of the NDCGs were associated with highly specific gene ontology (GO) terms (Fig. 3B and Table S3) and were named according to biological functions (either the top GO terms or function identified after literature review). The following NDCGs (Fig. 3A-C, Table S2-3) were identified: “Ribosome” was composed almost entirely of ribosome subunit genes. “Insulin secretion” contained multiple key genes for insulin secretion, including *G6PC2, ABCC8, SLC30A8, KCNK16, PCSK1*, and *PCSK2*. “Lysosome” was adjacent to “Insulin secretion” and contained genes related to lysosomal function, e.g. *GAA, PSAP, CTSA* and also genes with roles in regulation of insulin secretion, e.g. *SYT7, CD59, KIAA1324*. Unfolded protein response (“UPR”) contained stress-related genes including *TRIB3, DDIT3, DDIT4, EIF4EBP1*, and *EIF2S2*. Also, *ARG2*, previously shown to be affected in T2D was present^5^. “Microfilaments” contained genes related to actin organization e.g. *ACTN1, MARCKS*, and *WASL*. “Glycolysis” contained the key regulators of glycolysis, *ENO1, ALDOC, PGAM1*, and *TPI1*. “Proliferation” contained *PAX6* and multiple genes involved in cell proliferation and differentiation. Also, *ISL1* was present further up in this network. “Glucose response” contained a high concentration of genes crucial for beta cell function, e.g. *PDX1, NEUROD1, MAFA, MAFB, GCK*, and *FFAR1*. “Microtubuli” contained genes regulating microtubules and *NKX2*.*2*. “Mitochondria” was dominated by genes encoding subunits of complex 1 and 4 of the electron transport chain of the mitochondria. “Cell cycle” contained *STX5*, an important regulator of insulin secretion.

**Figure 3.**
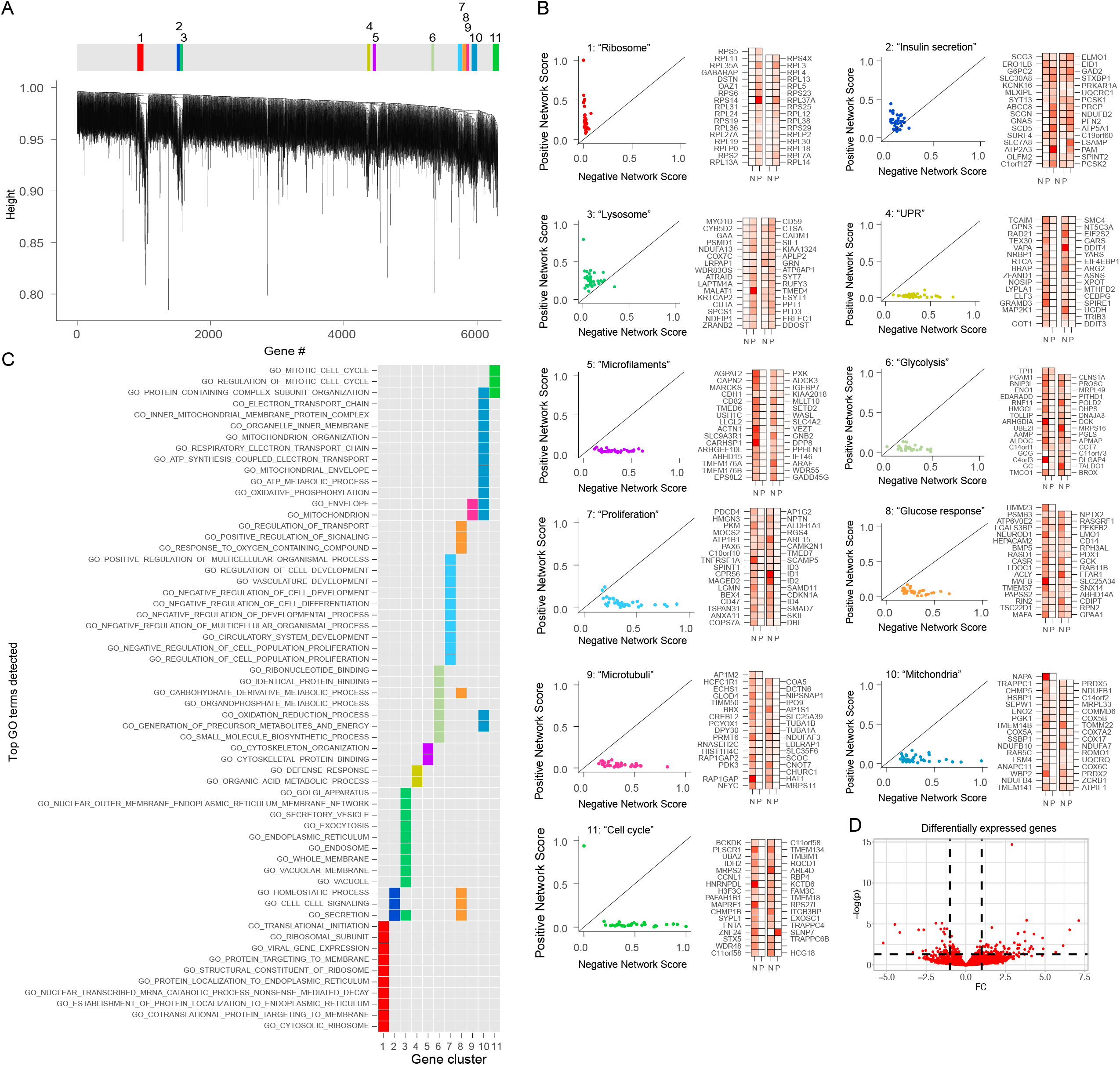
dGCNA uncovers the functional context of genes that correspond to canonical and non-canonical T2D pathways of beta cells. Dendrogram and clustering of 11 NDCGs in beta cells **(A)**. Scatter plots representing coordination rank scores for genes in each NDCG with degree of negative and positive scores indicated in red **(B)**. GO-term (biological process) enrichment is highly specific for each NDCG **(C)**. Columns represent NDCGs and rows are top GO-terms.

To investigate if the NDCGs were either hyper- or de-coordinated we assigned each gene a rank score depending on the eigen-vector centrality (dependent on the number of connections with other genes that was lost or gained) (Fig. 3B). Three NDCGs exhibited a general hyper-coordination in T2D (“Ribosome”, “Insulin secretion”, and “Lysosome”) and eight were overall de-coordinated (“UPR”, “Microfilaments”, “Glycolysis”, “Proliferation”, “Glucose response”, “Microtubuli”, “Mitochondria”, and “Cell cycle”). Plotting the top enriched GO-terms (Table S3) for each NDCG across all NDCGs (Fig. 3C) revealed a high degree of specificity in terms of biological annotation of the identified NDCGs. Thus, dGCNA, without prior information, identified specific biological processes including all established T2D dysfunction pathways in beta cells, but also genes and processes not previously associated with beta cell dysfunction. Of note, the expression levels of individual genes or clusters of genes did not need to change to be included in NDCGs. Differentially expressed genes are presented in (Fig 3D; Table S4).

We next used the two independent data sets (Dataset 1 and 2) to test if our findings replicated across datasets. Using dGCNA we observed 26% of overlap in genes within networks with altered coordination in T2D beta cells (Fig. 4A). Furthermore, comparing GO-terms of the genes from networks with altered coordination (Table S3) revealed a striking degree of overlap between the two data sets (Fig 4B). To compare overlaps between data sets using dGCNA as compared with differential expression on data averaged across each cell type in donors (pseudo-bulk DESeq2, adjusted for age, sex and BMI; Table S4) for the two data sets. We observed very few significant differentially expressed genes using DESeq2 (Table S4). To compare the methods, we thus ranked differentially expressed genes based on their p-value and compared their overlap up to the number of genes indicated by dGCNA (Fig. 4C). We observed substantially greater overlap using dGCNA as compared to DESeq2. In terms of reproducibility, dGCNA thus outperforms differential expression analysis also between data sets.

**Figure 4.**
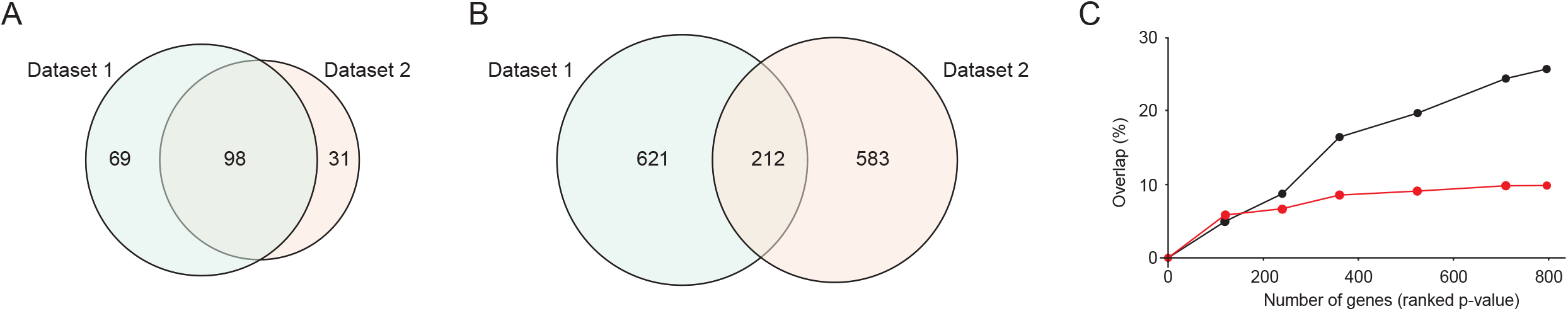
dGCNA outperforms DESeq2 for replication of T2D-affected genes between data sets. Overlap of genes identified with dGCNA in data set 1 and 2 **(A)**. Overlap of GO-terms on dGCNA genes for data set 1 and 2 **(B)**. Comparison of overlap of genes between data set 1 and 2 using DESeq2 and dGCNA on top ranked genes 50-800 genes **(C)**.

To test the predictive power of dGCNA in identifying genes involved in T2D-related processes we selected seven genes without a well-known presence or function in beta cells that were highly ranked in five of the NDCGs (Fig. 5). We confirmed the presence of the protein products in human pancreatic sections (Fig. 6A,H) and Fig S2) and performed siRNA silencing (two for each gene) of gene homologues in rat INS-1 832/13 cells (Fig. S3A). Silencing each of the seven genes resulted in increased mRNA expression of *Ins1* or *Ins2* (Fig. 5 A,B) and Fig. S3B-G). Furthermore, silencing five of the selected genes resulted in affected insulin secretion stimulated with glucose or glucose and IBMX^26^ (Fig. 5C,D). The recently described regulator of beta cell insulin-sensitivity inceptor/*KIAA1324*^*27*^ was highly ranked in (“Lysosome”) and its protein expression was highly increased in beta cells in pancreatic sections from T2D donors (Fig 5E,F). Furthermore, in agreement with the two identified NDCGs related to cytoskeletal organization (“Microfilaments” and “Microtubuli”), we confirmed altered cytoskeletal organization in beta cells in pancreatic sections from T2D donors (Fig 5G,H).

**Figure 5.**
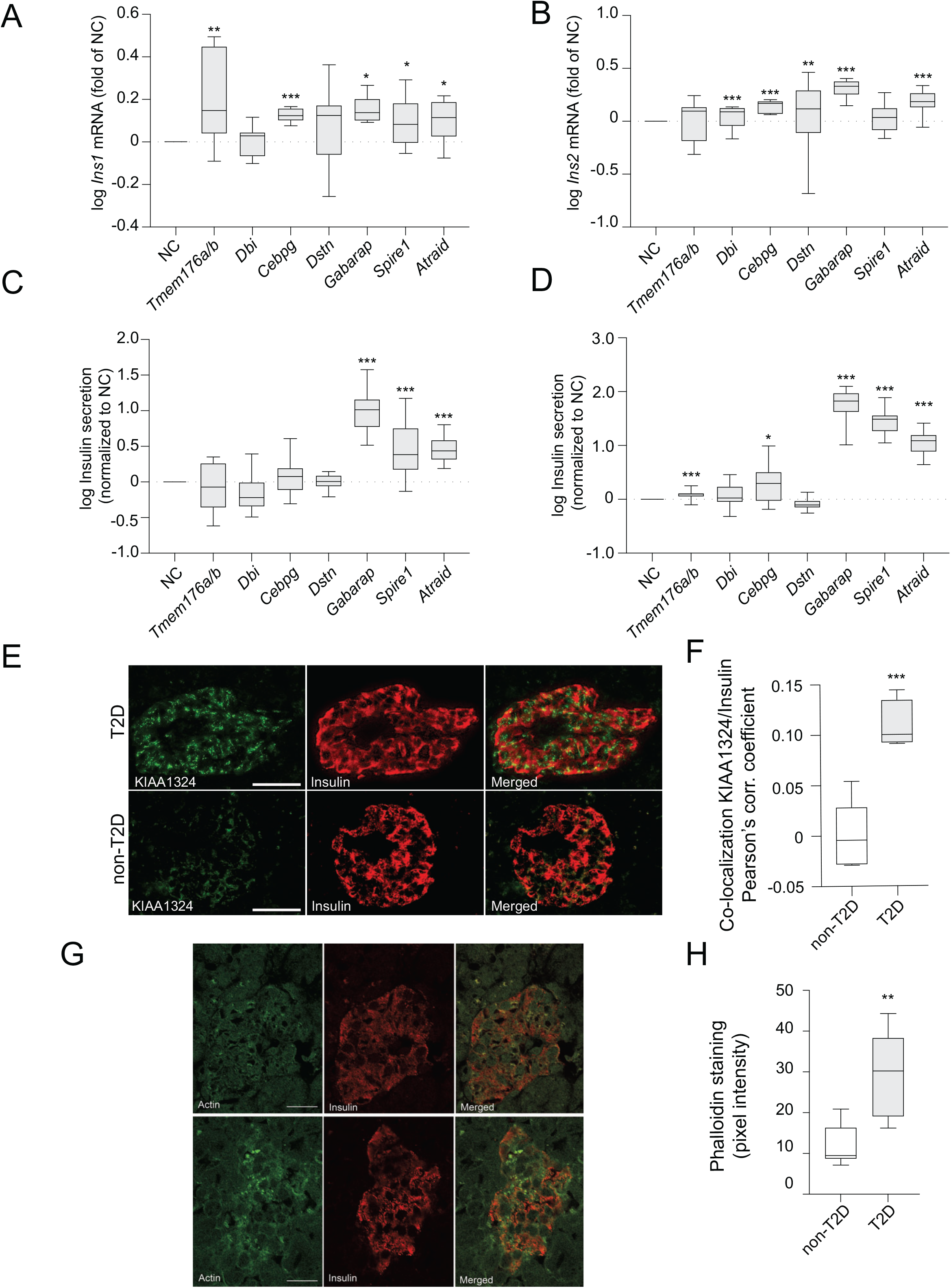
Effect of target genes on *Ins1* and *Ins2* mRNA, and on glucose-stimulated insulin secretion. siRNA silencing of *Tmem176a/b, Cebpg, Gabarap, Spire1* or *Atraid* resulted in increased *Ins1* mRNA levels **(A)** and silencing of *Dbi, Cebpg, Dstn, Gabarap* or *Atraid* increased *Ins2* mRNA **(B)** in INS-1 832/13 cells. Silencing of *Gabarap, Spire1* or *Atraid* caused increased insulin secretion in INS-1 832/13 cells at 16.7 mM glucose **(C)**. At 16.7mM glucose with IBMX silencing of *Tmem176a/b, Cebpg, Gabarap, Spire1* or *Atraid* resulted in increased insulin secretion **(D)**. T2D donors have increased beta cell immunoreactivity for KIAA1324 compared with non-T2D donors **(E)**. Quantification of colocalization of insulin and KIAA1324 in non-T2D (n=6) and T2D donors (n=4) **(F)**. T2D donors have more actin in beta cells compared with non-T2D donors **(G)**, quantified as pixel intensity in **(H)** in non-T2D (n=8) and T2D donors (n=5). Scale bar in E and G is 50 µm. See also Fig. S3.

Next, we used two knockout mouse models for high-ranked genes without prior association with beta cell function or T2D. *TMEM176A/B* (“Microfilaments”) are a pair of duplicated genes encoding non-selective cation channels^28^ reported to regulate cancer growth^29^. On a normal diet, *Tmem*176*a/b* knockout mice^30^ were normoglycemic and showed unaffected insulin- and glucose responses during an intraperitoneal glucose tolerance (Fig. S4A-D). In contrast, *Tmem*176*a/b* knockout mice exhibited reduced beta cell mass (Fig. 6B) and islet size (Fig. 6C), in line with the reported role in cell growth. On a high fat diet *Tmem*176*a/b* knockout mice exhibited a negative acute insulin response (Fig. 6D, Fig. S4E) and perturbed glucose elimination (Fig. 6E) during an intraperitoneal glucose tolerance test. Furthermore, *Tmem*176*a/b* knockout mice had altered cytoskeletal organization (Fig. 6F,G).

**Figure 6.**
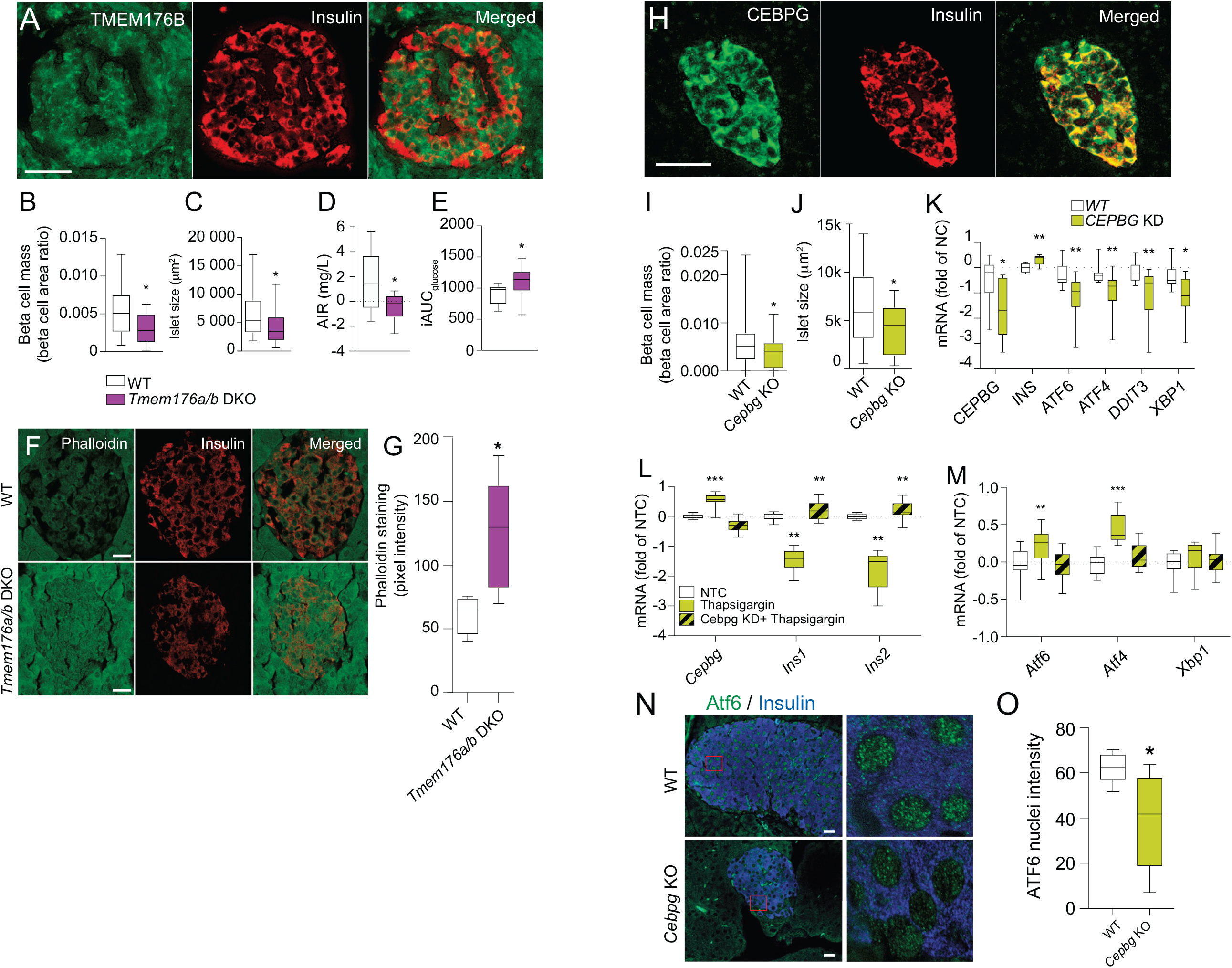
The function of hub genes *TMEM176A/B* and *CEBPG* in beta cells correspond to the context revealed by dGCNA. *TMEM176B* protein is expressed in human beta cells **(A)**. Reduced beta cell mass (**B**) and islet size (**C**) in *Tmem176a/b* double knockout (DKO) mice. Reduced acute insulin response (AIR) (**D**) and higher glucose levels calculated as iAUC (**E**) during IpGTT in *Tmem176a/b* DKO mice fed a high-fat diet. **(F and G)** *Tmem176a/b* DKO mice show approximately 2-fold increased actin staining in beta cells. *CEBPG* protein is expressed in human beta cells **(H)**. Reduced beta cell mass (**I**) and islet size (**J**) in *Cebpg* KO mice. **(H)** *CEBPG* silencing in human islets reduces the expression of *ATF6, ATF4, XBP1* and *DDIT3* **(H)**, all key mediators of UPR. Increased expression of *Cebpg* and decreased expression of *Ins1* and *Ins2* in thapsigargin-treated INS-1 832/13 cells **(L)**. Conversely, *Ins1* and *Ins2* expression was increased by thapsigargin treatment after *Cebpg-*silencing. The increased expression of *Atf6* and *Atf4* upon thapsigargin treatment was abolished after *Cebpg* silencing **(M)**. NTC, non-treated cells. *Cebgp* KO mice have reduced nuclear expression of ATF6 **(N and O)**. n=6 per group. Scale bar=50 µm. See also Fig. S4.

*CEBPG* (“UPR”) controls redox homeostasis in normal and cancerous cells^31^. *Cebpg* knockout mice were normoglycemic but had lower beta cell mass and smaller islets compared with WT controls (Fig. 6I,J). To test the dGCNA-prediction that *CEBPG* is central in the UPR pathway we silenced *CEBPG* in human islets resulting in reduced mRNA expression of *ATF4, ATF6*, and *XBP1*, all key mediators of UPR (Fig. 6K). As in INS-1 cells, *CEBPG* KD caused increased *INS* expression. Furthermore, thapsigargin (UPR-inducer) increased expression of *Cebpg* and decreased *Ins1* and *Ins2* mRNA. Conversely, *Ins1* and *Ins2* mRNA increased upon thapsigargin treatment when *Cebpg* was silenced (Fig. 6L). Similarly, thapsigargin increased *Atf4* and *Atf6* mRNA expression but failed to do so when *Cebpg* was silenced (Fig. 6M). Finally, *Cebpg* KO mice had less nuclear expression of ATF6 in beta cells (Fig. 6N,O). Our results thus confirm that *CEBPG* is a key-regulator of beta cell UPR and that *TMEM176A/B* are involved in the control of beta cell cytoskeleton arrangement.

To examine if T2D is associated with changed coordination in other islet cell types we also analyzed alpha cells (Fig. 7). We identified four NDCGs in alpha cells (Fig. 7A) and named them according to biological functions (top GO terms or literature review) (Fig. 7B and, Table S2, and S5). “Secretory granules” (hyper-coordinated) had GO-terms related to secretory granules, and contained *CHGB* and *SCG2*, as well as *CPE* and *PAM* (important for peptide hormone activation) and several genes related to protein folding. “Glycolysis” (de-coordinated) contained multiple genes with key roles in glycolysis or hexose metabolism, including *ALDOA, TPI1, PGK1, GAPDH, ENO1, ENO2, LDHA*. “Mitochondria” (de-coordinated) contained *SNAP25, CALM2 VDAC1, VDAC2*, and *SRI*, suggesting a role also in Ca^2+^ signaling. “Ribosome” (de-coordinated) had strong GO terms related to translation and contained multiple ribosome subunit genes (Fig. 7C, Table S2, and S5). Again, GO-term analysis revealed a notable specificity for each NDCG (Fig. 7B). Also for alpha cells there was a significant overlap between data set 1 and data set 2 for genes identified with dGSNA (Fig. 7E) as well as for the GO-terms (Fig. 7D).

**Figure 7.**
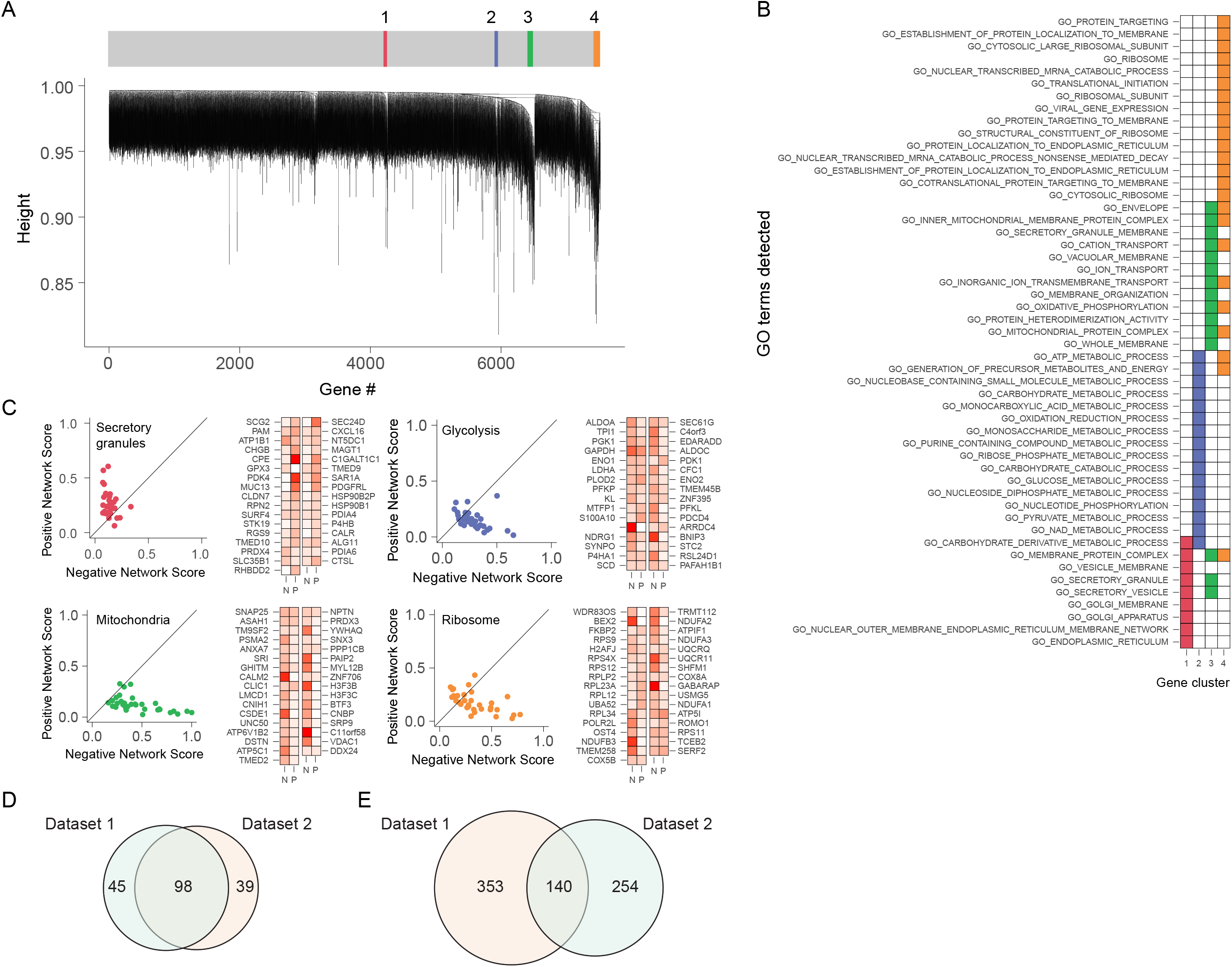
dGCNA uncovers non-canonical T2D NDCGs in alpha cells. Dendrogram of 4 NDCGs in alpha cells **(A)**. GO-term (biological process) enrichment is highly specific for each NDCG **(B)**. Columns represent NDSGs and rows are top GO-terms. Scatter plots representing coordination rank scores for genes in each NDSG and genes at the tip of each NDCG with negative and positive score indicated **(C)**. Venn diagram of the overlap of GO-terms detected by dGCNA in alpha cells between data set 1 and data set 2 **(D)**. Venn diagram of the overlap of genes in alpha cells between data set 1 and data set 2 **(E)**.

## DISCUSSION

Here we introduce dGCNA as a tool to interrogate scRNAseq data to reveal T2D-associated differential coordination of gene expression in islet cells, thereby inferring altered cellular functions. In addition to pathways and genes experimentally shown to underlie beta cell dysfunction, we identified non-canonical pathways and multiple genes with unknown roles in beta cell function. The ability of dGCNA to predict the functional context of T2D-affected beta cell genes was validated in both *in vitro* experiments and mouse models. Furthermore, we replicated our findings between two data sets and showed that dGCNA outperforms differential expression analysis for identification of T2D-associated changes in gene expression across datasets.

Using dGCNA we identified eleven NDCGs affected in T2D beta cells (Fig. 8). Most NDCGs contained multiple genes with established roles in beta cell function, as well as a multitude of genes not previously associated with T2D. A majority of the NDCGs were de-coordinated which we interpret as a perturbed function. In line with a body of evidence^32,33^, we observed mitochondrial dysregulation indicated by a de-coordinated NDCG containing genes encoding for Complex 1 and 4 of the electron transport chain. Also, in support of perturbed coordination of stimulus secretion coupling, we identified a de-coordinated NDCG containing key regulators of glycolysis and a de-coordinated NDCG (“Glucose response”) with a high density of key regulators of beta cell function, including *GCK* (the rate-limiting enzyme in glucose metabolism) suggesting perturbed glucose sensing in T2D beta cells.

**Figure 8.**
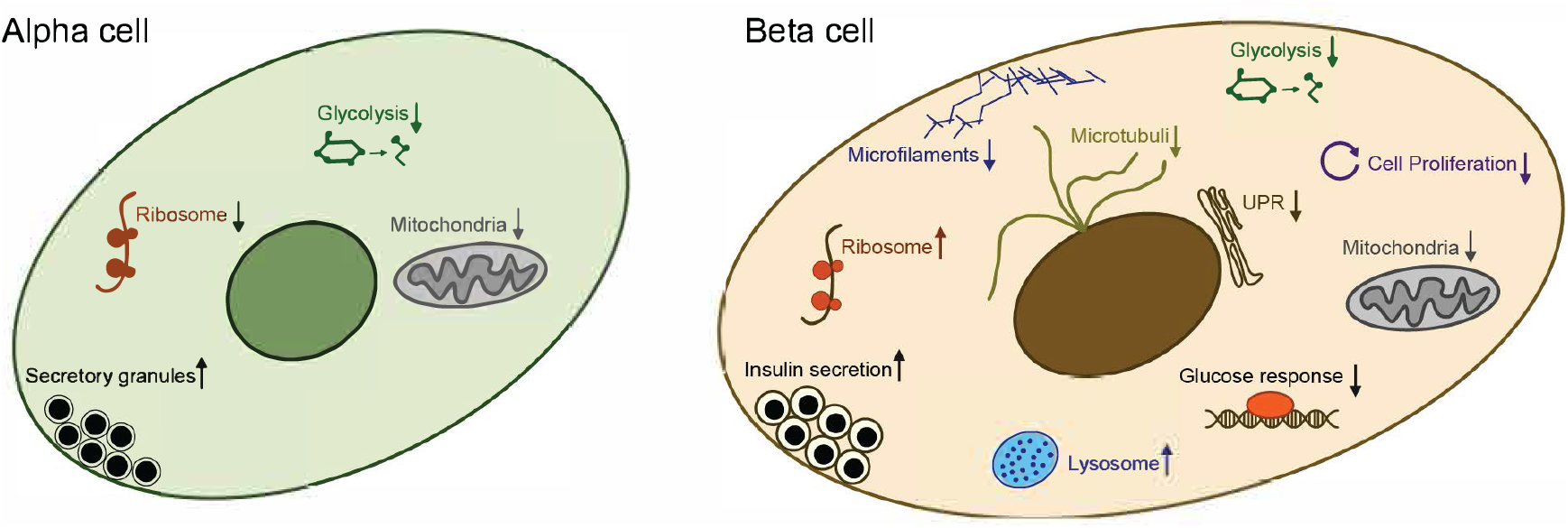
Biological processes affected in T2D alpha and beta cells. Hyper-coordinated processes marked with ↑ and de-coordinated processes marked with ↓.

Two NDCGs (“Proliferation” and “Microtubuli”) adjacent to “Glucose response” were also de-coordinated. These three NDCGs together contained virtually all key beta cell transcription factors, suggesting a general perturbation of beta cell transcriptional identity. “Proliferation” contained *PAX6, ISL1* and a number of genes known to regulate proliferation. Beta cell mass is controlled by a balance of apoptosis and renewal of beta cells^34^. Most studies agree that beta cell mass is reduced in human T2D, although its contribution to insulin deficiency has been questioned^1,35,36^. Our data is in support of reduced beta cell proliferation in T2D. Furthermore, “Microtubuli” containing *NKX2-2* and genes related to regulation of microtubules was de-coordinated. In further support of altered cytoskeletal arrangement in T2D beta cells, “Microfilaments”, containing multiple genes related to actin filament arrangement, was also de-coordinated. The role of actin in relation to insulin secretion is two-fold. Firstly, it acts as a physical barrier impeding the access of insulin granules to the cell periphery. Secondly, through glucose-induced remodeling it acts as a cytoskeletal structure for the transport of insulin granules to the plasma membrane^37,38^. Perturbed actin organization in T2D beta cells was confirmed in human pancreatic sections. Furthermore, *TMEM176A/B* in this network was identified as regulators of insulin production and secretion in cell lines and in a mouse model. Confirming a role for *TMEM176A/B* in regulation of cytoskeletal arrangement, *Tmem176a/b* KO mice had altered actin organization, reminiscent of that in T2D donors, and in agreement with the observed hampered insulin secretory response. Providing a plausible mechanistic link, *TMEM176* genes were recently shown to regulate Akt/mTOR-dependent pathways that are important regulators of actin remodulation^39^. Finally, we found a de-coordinated network of UPR genes. This indicates a perturbed capacity to remove misfolded proteins that could be contributing to beta cell failure in T2D. A central node gene in “UPR’’ was *CEBPG*, a redox-regulator that heterodimerizes with *ATF4*^*31*^. Our functional validation in a mouse model, cell lines and human islets put forward *CEPBG* as an important regulator of two out of three arms of the UPR pathway in beta cells.

These above-described processes likely represent pathophysiological mechanisms of T2D in beta cells. Conversely, we observed hyper-coordination in three NDCGs. “Ribosomes’’ was composed of ribosome subunit genes, indicative of increased translation of insulin. “Insulin secretion”, was composed of genes related to the exocytosis machinery, insulin processing and insulin granule content. Interestingly, a hyper-coordinated NDCG containing genes related to lysosomes was identified close to “Insulin secretion”. Lysosomes degrade insulin granules, via autophagy or crinophagy, and play an important role in protecting the beta cell in situations of stress^40^. The observation that this NDCG was devoid of autophagy genes speaks in favor of upregulated lysosomal degradation of insulin granules via crinophagy in T2D beta cells. The Lysosome NDCG also contained the lysosome-associated gene Inceptor/*KIAA1324*, recently identified in mice and human cell lines as a regulator of beta cell insulin sensitivity, confirming its altered regulation also in T2D patients^27^.

We also performed dGCNA on alpha cell data and identified four NDCGs affected in T2D alpha cells (Fig. 8). T2D alpha cells displayed a de-coordinated network of glycolysis genes. Glucagon secretion is tightly controlled by glucose and perturbed glycolysis is compatible with hampered control of glucagon secretion, a hallmark of T2D^41^. Contrary to beta cells, in T2D alpha cells a de-coordinated network composed of ribosome subunit genes was evident. This suggests reduced glucagon translation and could be interpreted as a compensatory mechanism, trying to lower glucagon secretion and thereby glucose output from the liver. Furthermore, a hyper-coordinated network with genes related to secretory granules and peptide hormone processing was evident.

Thus, by harnessing the individual variation between cells within a cell type the dGCNA algorithm identified cell type-specific, coordinated regulatory changes in T2D with unprecedented power. Our analysis generated an atlas of molecular changes in distinct populations of islet cells and thus allows for a bird’s-eye perspective of disease-related changes. Our findings support a model where basal insulin production and capability of exocytosis are enhanced in T2D beta cells, but the stimulus-secretion-coupling of insulin release and degradation of peptides that failed quality control are perturbed. This is in line with current clinical interventions based on enhancing insulin secretion using e.g. Sulfonylurea or GLP-1-based approaches^42^. In addition, our findings suggest that T2D beta cells are less differentiated, less capable of adaptive growth, display cytoskeletal remodeling compatible with perturbed insulin secretion, and signs of enhanced lysosome activity. Furthermore, our data point at cell type-specific alterations with non-overlapping programs in beta- and alpha cells.

While we assumed cell-to-cell variability, the coordination of biological programs reported herein are unbiased: we have listed all networks detected, each of which was highly specific with respect to GO-terms, supporting the hypothesis that the changes in gene-regulation were indeed coordinated on the level of pathways. Our analysis, without given prior information, revealed canonical pathways and multiple genes with experimentally proven importance in T2D beta cell dysfunction. It also revealed non-canonical processes and multiple genes not previously associated with T2D pathophysiology that could be potential therapeutic targets. The increased power of our analysis could potentially have great impact on the future design of perturbation experiments as well as clinical studies. Although we have not measured a temporal aspect of this coordination, we hypothesize that it represents changes in synchronization of gene expression. Interestingly, such a theory would entail that separate biological pathways are controlled with temporal specificity in cells in general. Our approach holds promise for use in elucidation of disease mechanisms of other complex tissues and diseases and warrants studies of the possibility of temporal coordination of gene expression.

## DATA AVAILABILITY

All data will be available on GEO upon publication. Data will, upon publication of the manuscript, be made available with correct sample identification on European Genome-phenome Archive (EGA).

Code availability: all code will be made public on Github upon publication.

## ACKNOWLEDGEMENTS

The authors would like to thank the Eukaryotic Single Cell Genomics facility at SciLifeLab where scRNAseq was performed, Human Tissue Lab at Lund University Diabetes Centre and The Nordic Network for Islet transplantation for providing islets. Rickard Fred for technical assistance, Karen Saylor and Nancy Martin for animal husbandry and genotyping. The authors are grateful to Claes Wollheim, Patrick F. Sullivan, Patrik Ernfors and Sten Linnarson for critical reading of the manuscript. This work was funded by joint grants to J.H-L and N.W. from the Novo Nordisk Foundation (grants NNF15OC0016546 and NNF16OC0021200). N.W. was supported by: the Swedish Research Council (2020-01017, 2017-00862 and 521-2012-2119), The Regional research foundation (ALF), Novo Nordisk Foundation (NNF20OC0063916), Diabetes Wellness Research Foundation Sweden, EFSD, The Hjelt Foundation, The Royal Physiographic Society in Lund, and The Swedish Diabetes Foundation. The Swedish Research Council (Dnr 2017-00862, Dnr 521-2012-2119, Linnaeus grant, Dnr 349-2006-237, and Strategic Research Area Exodiab, Dnr 2009-1039), the Swedish Foundation for Strategic Research (Dnr IRC15-0067). In addition, J.H-L. was supported by the Swedish Research Council (awards 2014-3863 and 2018-00799), the Wellcome Trust (108726/Z/15/Z); and ERC Grant Agreement [819540].

## MATERIALS AND METHODS

### Human subjects

Information about the donors included in the study is presented in Table S1. Human islets were obtained from the Nordic Network for Islet Transplantation (www.nordicislets.org). The ethics committees at Uppsala and Lund Universities approved all procedures.

### Animals

*Tmem176a/b* double knockout mice have been described in detail elsewhere^30^. For this study, the mice were kept in Macrolon cages in a temperature-controlled environment (21°C), on a 12h light/dark cycle and with free access to diet (standard rodent chow or high fat diet (60% fat, 20% protein, 20% carbohydrates, Special Diet Services, Witham, UK)) and tap water. All experiments were performed in accordance with institutional guidelines for animal care and use for the University of Nantes. The *Cebpg* KO mice^31^ were kept in Macrolon cages in a temperature-controlled environment (21°C), on a 12 h light/dark cycle and with free access to standard rodent chow and tap water. All animal experiments were approved by the animal ethics committee in Malmö and Lund, Sweden.

### Sample processing and single-cell RNA sequencing

Human islets were stored in 30 ml HEPES-buffered media. Islets were transferred to a 50 ml tube and centrifuged at 150 g for 2 min at RT. Supernatant was removed and cells were washed once with 5 ml of Accutase. Another 5 ml of Accutase was added for incubation at 37 °C for 8-10 min with pipetting up and down every other minute. After incubation, 5 ml of cold islet media was added and cells were suspended by pipetting. Cells were passed through a 40-μm cell strainer and centrifuged at 500 g for 5 min. Cells were washed twice with PBS at 500 g for 5 min. Finally, cells were resuspended in HBSS and kept on ice before FACS-sorting. Single-cells were FACS-sorted into 384-well plates. cDNA libraries were generated using the Smart-seq2 protocol^25^ and sequenced on a HiSeq 2500 sequencer.

### Filtering and cell type clustering

For data set 1. STAR 2.4.2a was used for alignments with the reference genome hg38 using 2-pass alignment for improved performance of *de nov*o splice junction reads, filtered for uniquely mapping reads only and saved in .bam files. The count matrix showed the individual counts aligning to each gene per cell. Expression values were computed as reads per kilobase of the gene model and million mappable reads (RPKMs). Sequenced cells with mRNA reads <100 000, percent of uniquely mapping reads <50 %, or percent of uniquely exonic reads <40 % were removed, obtaining 3645 transcriptomes. For dataset 2, raw sequencing data of human pancreatic islets were de-multiplexed and converted into fastq files using Illumina bcl2fastq with default setting. Reads from human data were aligned to human genome hg38 using the STAR aligner. Uniquely aligned reads mapping to the RefSeq gene annotations were used for gene expression estimation at reads per kilobase transcript and million mapped reads (RPKMs) using rpkmforgenes. Low quality cells were excluded from downstream analysis when they failed to meet the following criteria for retaining cells: 1) >= 50,000 sequence reads; 2) >= 40% of reads uniquely aligned to the genome; 3) >= 40% of these reads mapping to RefSeq annotated exons; and 4) >= 1000 genes with RPKM >= 1. In addition, doublets detected by Scrublet were further removed, resulting in a total of 4866 single cells for downstream analysis. For cell type assignment, top 1000 most variable genes were used for PCA dimension reduction followed by clustering using affinity propagation, resulting in seven exocrine cell clusters (acinar, ductal, PSC, MHC II, mast, stem like, endothelial cells) and one endocrine cell cluster. To further annotate endocrine cell types, we applied the same approach on endocrine cells only and identified five clusters representing alpha cells, beta cells, delta cells, gamma cells and epsilon cells, respectively.

For datasets 1 and 2, the expression of transcript variants with the same gene name were summed. The datasets were integrated and clustered using Conos and cell types were assigned using label propagation based on previous clustering analysis and known markers.

### Differential Gene Coordination Network Analysis (dGCNA)

Cell type-specific networks for each cell type with more than 30 cells/donor were constructed separately for cases and controls using pearson correlation between transcriptional levels in log2(RPKM+1) of each gene pair. We filtered non-expressed genes (RPKM+1 < 2) and genes detected (RPKM+1 > 2) in less than 5 cells/donor. The correlations were calculated using a linear mixed-effects model including random effects to correct for donor specific expression differences. To build cell type-specific differential networks between healthy and disease states we calculated the difference in pairwise Pearson correlations for each gene pair between controls and T2D cases. In order to remove false positive weights in the differential network, we used a bootstrapping approach generating random differential networks by the selection of 512 pairs of random groups of donors. We filtered all correlations with a relative frequency lower than 0.975 keeping only links that are significantly stronger than random. This approach provides an internally scaled thresholding that takes the quality of the underlying dataset into account. We then performed clustering of the differential networks (one per cell type) using *TOMdist (*WGCNA package) and hierarchical clustering (*hclust*) to reveal gene communities altered between the conditions. Networks of differentially coordinated genes (NDCGs), i.e. gene modules, were determined using the *cutreeDynamic* function (WGCNA package) at different threshold and were manually selected. Next, we evaluated whether these T2D-related NDCGs were associated with known biological functions using a hypergeometric test (Bonferroni correction, p<0.05) with Gene Ontology terms (Cellular component, Biological function and Molecular function). Protein coding genes detected in our dataset for each cell type were used as the background geneset. Network scores were calculated with the function *eigen_centrality* (igraph R package) for the positive (hyper coordinated) subnetwork (network with just positive weights) and for the negative (de-coordinated) subnetwork (network with just negative weights) separately.

### Differential expression analysis

We used the DESeq2 method to calculate differential expression of genes. Expression of all single-cell counts of the same cell type and the same donor were summed before calculating differential expression. We removed unwanted variation from the RNAseq counts with the package RUVSeq using control genes. Thresholds of the volcano plots were established to a fold change of 2 and an adjusted p-value of 0.5.

### INS-1 832/13 cell culture and siRNA-mediated gene silencing

INS-1 832/13 cells (*13*) were cultured at 5% CO_2_ and 37°C in RPMI1640 medium (Sigma Aldrich, St Louis, MO) containing 2 g/L D-glucose, supplemented with 10% fetal bovine serum, 10 mM HEPES, 1 mM sodium pyruvate and 50 μM β-mercaptoethanol (Sigma Aldrich).

Gene silencing in INS-1 832/13 cells was performed using Lipofectamine RNAiMAX (Life Technologies, Waltham, MA) and 60 nM siRNA targeting *Atraid* mRNA (s151757), *Cebpg* mRNA (s129415), *Dbi* mRNA (s128869), *Dstn* mRNA (282408), *Gabarap mRNA (s133040), Spire1* (s158071), *Tmem176a* mRNA (s150574) and *Tmem176b* mRNA (s140062). The sequences for scrambled siRNA were sense: 5’-GAGACCCUAUCCGUGAUUAtt-3’ and antisense: 5’- UAAUCACGGAUAGGGUCUCtt-3’ (Silencer Select, rat negative control #1; scrambled siRNA and all siRNAs were from Ambion, Life Technologies). Transfection complexes were prepared in accordance with the instructions provided by the manufacturer and added to 2×10^5^ cells seeded per well in 24-well plates.

Human islets were transfected with 60 nM siRNA targeting human *CEBPG* (s2901, Silencer Select Pre-designed siRNA, Ambion, Life Technologies), *TMEM176A* (s226845) or *TMEM176B* (s226234). The sequence for *CEBPG* siRNA were sense: 5’-AGAGCCGGUUGAAAAGCAAtt-3’ and antisense: 5’- UUGCUUUUCAACCGGCUCUtt-3’.The sequence for *TMEM176A* siRNA were sense: 5’- CUGAAGGCCUUGUUCAGAAtt-3’ and antisense: 5’-UUCUGAACAAGGCCUUCAGca-3’. The sequence for *TMEM176B* siRNA were sense: 5’-GAAGGAGGAGUGUAGAGCUtt-3’ and antisense: 5’-AGCUCUACACUCCUCCUUCtg-3’

### RNA extraction

Total RNA was isolated from INS-1 832/13 cells 48h after transfection using the NucleoSpin extraction kit (Macherey-Nagel, Düren, Germany). The amount of isolated RNA was measured using the NanoDrop system. For bulk RNA sequencing, samples were also analyzed using TapeStation. Total RNA from human islets was extracted using Nucleospin RNA XS (Macherey-Nagel) three days post-transfection.

### cDNA synthesis and real-time qPCR

For the experiment, 1 μg of isolated total RNA was reverse-transcribed to cDNA using the RevertAid First Strand cDNA synthesis kit (Life Technologies). qPCR was performed with 25 ng cDNA template using TaqMan Expression Master Mix (Life Technologies) according to the instructions provided by the manufacturer. *Tbp* and *Hprt1* (Rn01527840_m1 and Rn01455646_m1, respectively) were used as house-keeping genes. Expression levels were calculated using the 2^-ΔΔCt^-method. TaqMan assays used were: *Atf4* (Rn00824644_g1), *Atf6* (Rn01490844_m1), *Atraid* (Rn01468747_g1), *Cebpg* (Rn01764319_m1), *Dbi* (Rn00821402_g1), *Dstn* (Rn01415640_g1), *Gabarap* (Rn00490680_g1), *Ins1* (Rn0212433_g1), *Ins2* (Rn01774648_g1), *Spire1* (Rn01491339_m1), *Tmem176a* (Rn01451723_m1), *Tmem176b* (Rn00508100_m1) and *Xbp1* (Rn01443523_m1). For human islets, 300 ng of RNA was reverse transcribed to cDNA using RevertAid First Strand cDNA synthesis kit. RT-qPCR for target genes and two endogenous controls (*HPRT1* and *TBP*) was performed using 2.5 ng cDNA. TaqMans for human islet were: *ATF4* (Hs00909569_g1), *ATF6* (Hs00232586_m1), *CEBPG* (Hs01922818_s1), *DDIT3* (Hs99999172_m1), *HPRT1* (Hs4326321_m1), *INS* (Hs02741908_m1), *TBP* (Hs00427620_m1), *TMEM176A* (Hs00218506_m1), *TMEM176B* (Hs00962650_m1) and *XBP1* (Hs00231936_m1). All TaqMan assays were from Life Technologies and qPCR reactions were run on the Viia7 real-time PCR system (Applied Biosystems, Foster City, CA).

### Insulin secretion experiments

INS-1 832/13 cells were seeded in 24-well plates (2×10^5^ cells/well) and allowed to adhere for approximately 5 h before transfection. On the day of insulin secretion cells were washed in PBS, incubated with 2.8 mM glucose for 2 h and then incubated for 60 min with 2.8 mM glucose, 16.7 mM glucose or 16.7 mM glucose with 0.1 mM IBMX. For the experiments, glucose was dissolved in the secretion assay buffer (SAB; 114 mM NaCl, 4.7 mM KCl, 1.2 mM KH_2_PO_4_, 1.16 mM MgSO_4_, 1.16 mM CaCl_2_, 20 mM HEPES, 25.5 mM NaHCO_3_ and 0.2% BSA). The cells were then placed on ice and the culture media was collected for insulin and protein determinations. Insulin concentration was determined by ELISA from Mercodia (Uppsala, Sweden) and protein concentration was determined using Protein Assay Dye Reagent (BioRad, Hercules, CA).

### Thapsigargin treatment

INS-1 832/13 cells were seeded in 24-well plates (2×10^5^ cells). 100 nM thapsigargin (Sigma Aldrich) was added to the cells for 24 h. In a second experiment, cells were seeded in 24-well plates and *Cebpg* mRNA was silenced using siRNA (s129415) for 24 h. 100 nM thapsigargin was then added to these cells for 24 h. Total RNA was extracted from the cells from both above-mentioned experiments.

### Immunohistochemistry

Primary antibodies: anti-rabbit ATF6 (code NBP1-76675, dilution 1:200, Novus Biologicals, Centennial, CO), anti-rabbit CEBPG (code HPA012024, dilution 1:20, Sigma Aldrich), anti-rabbit DSTN (code ab186754, dilution 1:100, Abcam), anti-rabbit GABARAP (code HPA-78365, 1:500 dilution, Sigma Aldrich), anti-guinea pig insulin (code A0564, 1:1000 dilution, Dako, Glostrup, Denmark), anti-rabbit KIAA1324 (code PA5-67123, dilution 1:800, Life Technologies), anti-rabbit SPIRE1 (ab130403, 1:1000 dilution, Abcam), anti-rabbit TMEM176A (code NBP1-83283, 1:20 dilution, Novus Biologicals) and anti-rabbit TMEM176B (code CSB-PA023758LA01HU, 1:50 dilution, CusaBio, Houston, TX). All antibodies were diluted in PBS with 0.25% BSA and 0.25% Triton X-100. Slides were incubated with primary antibodies overnight at 4°C, then washed twice in PBS with 0.25% Triton X-100. Slides were then incubated for 1 h with secondary antibodies. Secondary antibodies used were: Donkey anti-guinea pig AlexaFluor594 (for insulin), donkey anti-rabbit Cy2 (for ATF6, CEBPG, DSTN, GABARAP, KIAA1324, SPIRE1, TMEM176A and TMEM176B) and goat anti-guinea pig AlexaFluor405 (for insulin). For phalloidin staining, Phalloidin-iFluor488 (Abcam) was diluted in PBS (+0.25% Triton X-100 and 0.25% BSA; 1:1000) and incubated for 60 min at room temperature. Slides were then washed twice in PBS with 0.25% Triton X-100 and mounted. Immunofluorescence was examined in an epi-fluorescence microscope (Olympus BX60, Olympus, Tokyo, Japan) or a Zeiss LSM800 confocal microscope (Oberkochen, Germany). Images were taken with a digital camera (Olympus DP74, Olympus) using the CellSens software (Olympus) or Zen System 2.6 (Zeiss). Pixel intensity was measured using ImageJ software and colocalization (expressed as Pearson’s colocalization coefficient) analysis was performed using the CellSens software.

### Beta cell mass measurements in *Cebpg* KO mice and *Tmem176a/b* DKO mice

Beta cell mass measurements in the KO mouse models were performed at three depths of the pancreas (100 μm difference between a series of sections). Insulin was visualized using an anti-insulin antibody (code A0564, 1:1000 dilution, Dako) and anti-guinea pig Alexa Fluor 594 (1:400 dilution) and the signal was used to determine the total islet area which was divided by the total pancreas area to generate a measure of beta cell mass. The average islet size and total islet number were also calculated. Immunofluorescence was examined under an epifluorescence microscope (Olympus BX60, Olympus, Tokyo, Japan) with a digital camera (Olympus DP74, Olympus). Measurements were made at 40X magnification using the CellSens Dimensions software (Olympus).

### Intraperitoneal glucose tolerance test (IpGTT)

After a 4-h fasting period, basal blood samples were drawn via retro-orbital puncture. The mice were then injected peritoneally with glucose (2 g/kg bodyweight). Blood samples were collected after 10, 20, 30, 60 and 90 minutes into chilled EDTA-tubes. Samples were maintained on ice and centrifuged at 500xg for 5 min and plasma stored at -80°C until analysis.

### Insulin secretion and glucose measurement

Plasma insulin from the *Tmem176a/b* DKO mice subjected to IpGTT and fasting plasma insulin from the *Cebpg* KO mice was determined by ELISA (Mercodia) according to the manufacturer’s instructions. Glucose from the mice was analyzed using the Infinity Glucose (Ox) kit from Thermo Scientific according to the instructions provided by the manufacturers. Acute insulin response (AIR) in the *Tmem176a/b* DKO mice was calculated at the difference between 10-min insulin levels and basal insulin levels. Integrated area under the curve (iAUC) for the glucose levels in the *Tmem176a/b* DKO mice subjected to IpGTT was calculated using GraphPad Prism 8 (GraphPad Prism, San Diego, CA).

## SUPPLEMENTARY MATERIALS

Figs. S1 to S4

Tables S1 to S5

## LEGENDS TO SUPPLEMENTARY FIGURES

**Suppl Fig 1 (Related to Fig 1)**.

**(A)** Number of detected genes per cell type. **(B)** UMAP of 8511 cells with donors indicated.

**Suppl Fig 2 (Related to Fig 6)**.

Double immunostaining for target gene protein products and insulin in human pancreatic sections shows expression of TMEM176A (top left panels), DSTN (top right panels), SPIRE1 (bottom left panels) and GABARAP (bottom right panels) in human beta cells. Scale bar=50 µm.

**Suppl Fig 3 (Related to Fig. 5). KD efficiency and individual pairs of siRNAs**. Knockdown efficiency of each pair of siRNAs used **(A)**. All pairs of siRNA significantly (p<0.05) reduced the expression of the gene of interest compared with scrambled control siRNA (dashed line). (**B-G**). Effect of each pair of siRNAs on *Ins1* and *Ins2* mRNA.

**Suppl Figure 4 (Related to Fig. 6). Extended *Tmem176a/b* DKO mouse data**.

No difference detected in *Tmem176a/b* DKO mice fed a normal diet during an intraperitoneal glucose tolerance test (IpGTT) with respect to insulin **(A)** and glucose **(B)** levels, as well as acute insulin response (AIR) **(C)** and iAUC for glucose **(D)**. *Tmem176a/b* DKO mice fed a high-fat diet had a negative insulin response (**E**), (AIR in **Fig. 6D**) and higher postprandial glucose levels (**F**), (iAUC in **Fig. 6E**) during an IpGTT.

