## Supplementary material for "Single-cell mRNA-regulation analysis reveals cell type-specific mechanisms of type 2 diabetes": Suppl figures and tables

Fig. S1

A

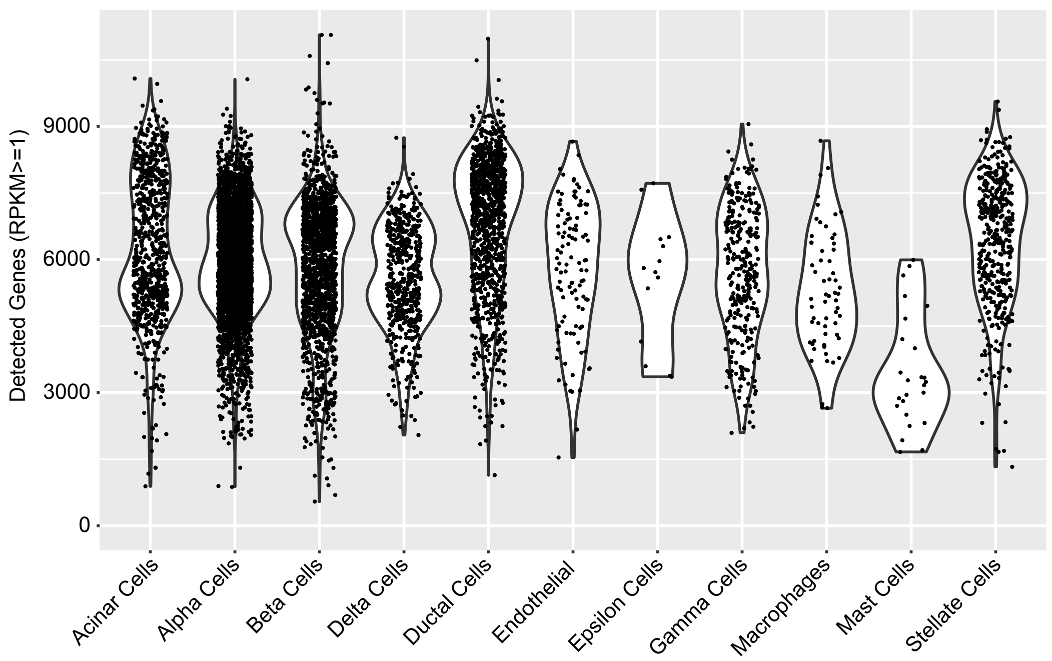

Detected Genes (RPKM>=1)
9000
6000
3000
0
Acinar Cells
Alpha Cells
Beta Cells
Delta Cells
Ductal Cells
Endothelial
Epsilon Cells
Gamma Cells
Macrophages
Mast Cells
Stellate Cells


B

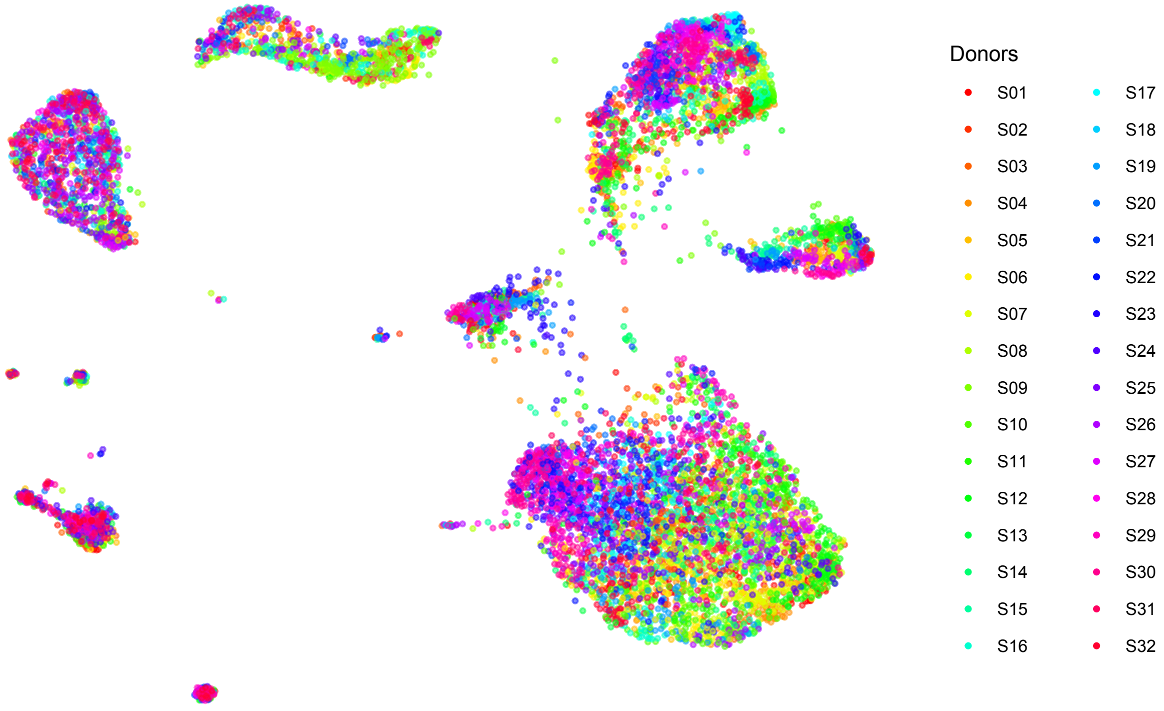

Donors
S01
S02
S03
S04
S05
S06
S07
S08
S09
S10
S11
S12
S13
S14
S15
S16
S17
S18
S19
S20
S21
S22
S23
S24
S25
S26
S27
S28
S29
S30
S31
S32

### Fig. S2

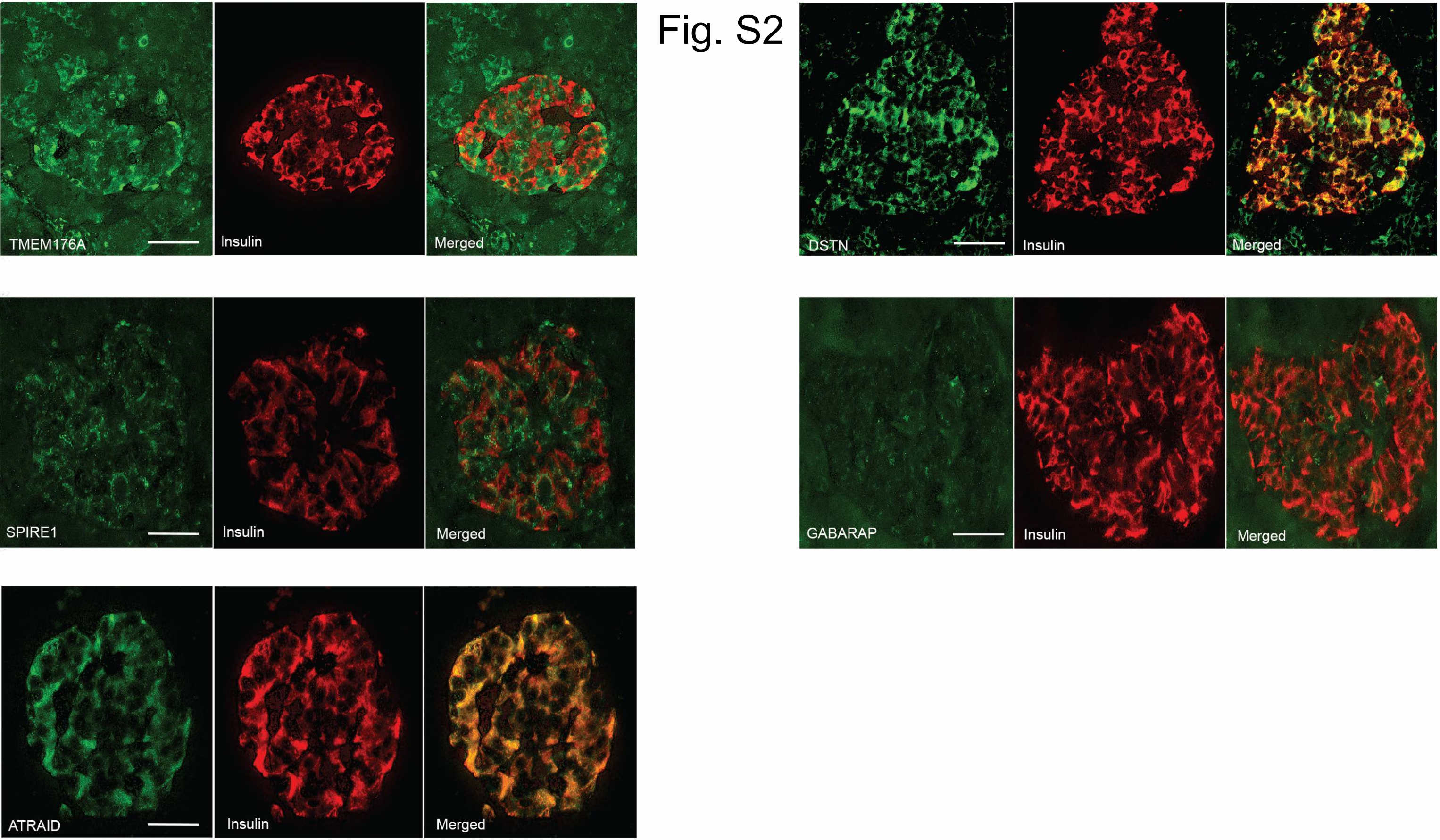
TMEM176A
Insulin
Merged
DSTN
Insulin
Merged
SPIRE1
Insulin
Merged
GABARAP
Insulin
Merged
ATRAID
Insulin
Merged

Fig. S3

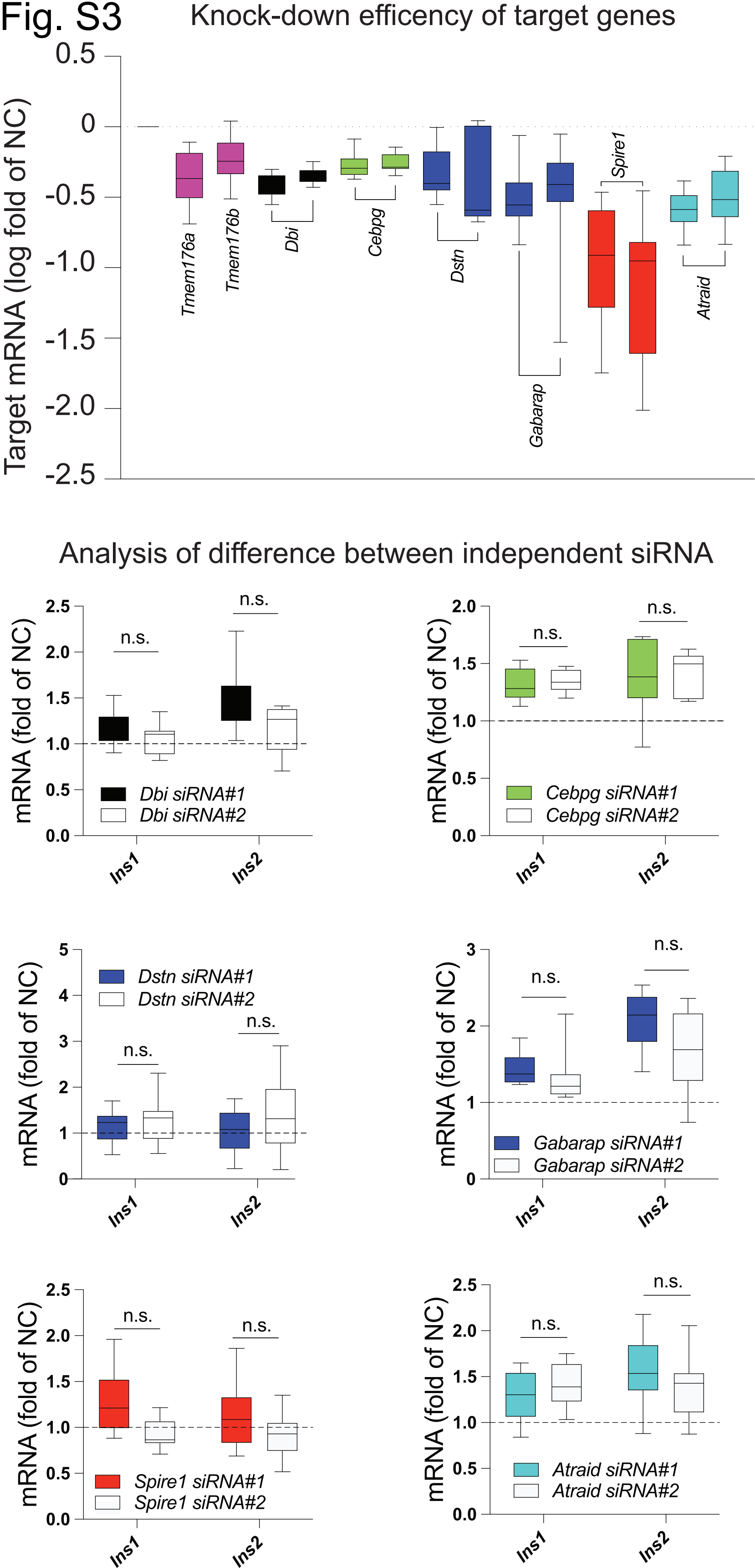
Knock-down efficency of target genes
Target mRNA (log fold of NC)
0.5
0
-0.5
-1.0
-1.5
-2.0
-2.5
Tmem176a
Tmem176b
Dbi
Cebpg
Dstn
Gabarap
Spire1
Atraid
Analysis of difference between independent siRNA
mRNA (fold of NC)
n.s.
Dbi siRNA#1
Dbi siRNA#2
Cebpg siRNA#1
Cebpg siRNA#2
Dstn siRNA#1
Dstn siRNA#2
Gabarap siRNA#1
Gabarap siRNA#2
Spire1 siRNA#1
Spire1 siRNA#2
Atraid siRNA#1
Atraid siRNA#2
Ins1
Ins2

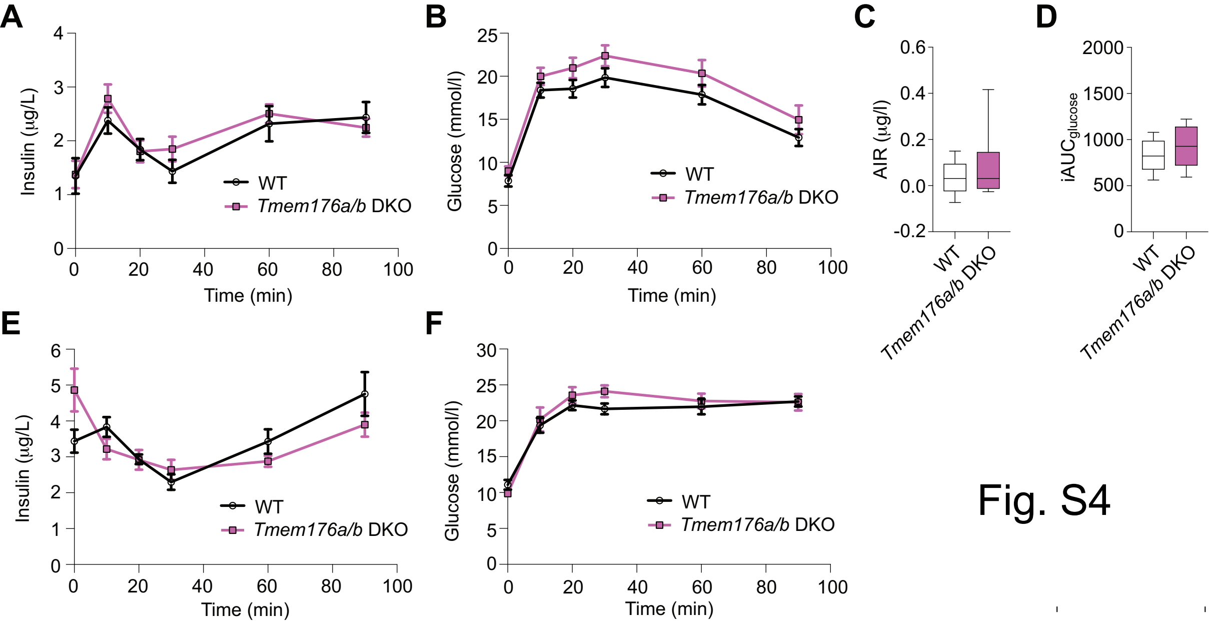
A
Insulin (μg/L)
Time (min)
WT
Tmem176a/b DKO
B
Glucose (mmol/l)
Time (min)
WT
Tmem176a/b DKO
C
AIR (μg/l)
WT
Tmem176a/b DKO
D
iAUCglucose
WT
Tmem176a/b DKO
E
Insulin (μg/L)
Time (min)
WT
Tmem176a/b DKO
F
Glucose (mmol/l)
Time (min)
WT
Tmem176a/b DKO

Fig. S4

**Table S1 Donor charachteristics**

| Sample ID | Donor ID | Sex | Age (years) | BMI (Kg/m2) | HbA1c (%) | T2D |
|---|---|---|---|---|---|---|
| S01 | 1 | Female | 65 | 20.8 | 5.9 | No |
| S02 | 2 | Male | 24 | 33.0 | 5.9 | No |
| S03 | 3 | Female | 58 | 24.4 | 5.9 | No |
| S04 | 4 | Male | 42 | 29.2 | 5.8 | No |
| S05 | 5 | Female | 73 | 22.3 | 5.8 | No |
| S06 | 6 | Male | 66 | 24.3 | 5.7 | No |
| S07 | 7 | Male | 64 | 34.0 | 7.9 | Yes |
| S08 | 8 | Male | 49 | 50.9 | 6.3 | Yes |
| S09 | 9 | Female | 59 | 29.4 | 6.6 | Yes |
| S10 | 10 | Female | 56 | 29.5 | 6.9 | Yes |
| S11 | 11 | Male | 58 | 26.8 | 6.0 | Yes |
| S12 | 12 | Male | 63 | 27.8 | 5.6 | Yes |
| S13 | H1 | Male | 43 | 30.8 | 5 | No |
| S14 | H10 | Female | 45 | 23.03 | 4.6 | No |
| S15 | H2 | Male | 25 | 24.67 | 5.3 | No |
| S16 | H3 | Female | 48 | 35.0 | 5.7 | No |
| S17 | H4 | Male | 22 | 32.95 | 5.4 | No |
| S18 | H5 | Male | 27 | 31.8 | N/A | No |
| S19 | H6 | Male | 23 | 21.45 | 5.5 | No |
| S20 | H7 | Female | 31 | 28.72 | 5.4 | No |
| S21 | H8 | Male | 39 | 25.38 | 5.5 | No |
| S22 | H9 | Female | 36 | 31 | 5.6 | No |
| S23 | T2D1 | Male | 57 | 23.98 | 7.0 | Yes |
| S24 | T2D10 | Male | 51 | 35.59 | 7.1 | Yes |
| S25 | T2D2 | Female | 37 | 39.6 | 7.3 | Yes |
| S26 | T2D3 | Male | 52 | 34.36 | 7.7 | Yes |
| S27 | T2D4 | Female | 55 | 29.84 | 7.4 | Yes |
| S28 | T2D5 | Female | 41 | 42.14 | 6.5 | Yes |
| S29 | T2D6 | Male | 48 | 43.7 | 6.6 | Yes |
| S30 | T2D7 | Male | 51 | 24.4 | 6.9 | Yes |
| S31 | T2D8 | Female | 42 | 27.6 | 6.7 | Yes |
| S32 | T2D9 | Male | 59 | 32.5 | 6.6 | Yes |

Table S2: NDCGs in beta cells.

| |
|---|
| **Branch 1** |
| RPS5 RPL11 RPL35A GABARAP DSTN OAZ1 RPS6 RPS14 RPL31 RPL24 RPS19 RPL36 RPL27A RPL19 RPLP0 RPS2 RPL13A RPS4X RPL3 RPL4 RPL13 RPL5 RPS23 RPL37A RPS25 RPL12 RPL38 RPS29 RPLP2 RPL30 RPL18 RPL7A RPL14 ATP5L FAU RPL37 RPL10A PFDN5 RPS20 RPL8 RPS7 RPS17 PEBP1 RPL36AL UBB EEF2 TPT1 UBL5 COX6B1 UQCR11 ATP5G2 COX8A LAMTOR4 CHMP2A SLIRP TCEB2 NDUFS5 |
| **Branch 2** |
| SCG3 ERO1LB G6PC2 SLC30A8 KCNK16 MLXIPL SYT13 ABCC8 SCGN GNAS SCD5 SURF4 SLC7A8 ATP2A3 OLFM2 C1orf127 ELMO1 EID1 GAD2 STXBP1 PRKAR1A UQCRC1 PCSK1 PRCP NDUFB2 PFN2 ATP5A1 C19orf60 LSAMP PAM SPINT2 PCSK2 |
| **Branch 3** |
| MYO1D CYB5D2 GAA PSMD1 NDUFA13 COX7C LRPAP1 WDR83OS ATRAID LAPTM4A MALAT1 KRTCAP2 CUTA SPCS1 NDFIP1 ZRANB2 PSAP CD59 CTSA CADM1 SIL1 KIAA1324 APLP2 GRN ATP6AP1 SYT7 RUFY3 TMED4 ESYT1 PPT1 PLD3 ERLEC1 DDOST DHRS7 DZIP3 ABCA5 OCIAD1 CLDN7 TAPBP PON2 COX6A1 TMEM205 |
| **Branch 4** |
| TCAIM GPN3 RAD21 TEX30 VAPA NRBP1 RTCA BRAP ZFAND1 NOSIP LYPLA1 ELF3 GRAMD3 MAP2K1 EPB41L4A-AS1 GOT1 CRCP SMC4 NT5C3A EIF2S2 GARS DDIT4 YARS EIF4EBP1 ARG2 ASNS XPOT MTHFD2 CEBPG SPIRE1 UGDH TRIB3 DDIT3 TCEA1 MIR6758 MARS PCK2 RNF187 RBCK1 MAP7 SYNCRIP THUMPD1 BCL7B ZFAND5 COG5 DNTTIP2 ZNF770 MBOAT7 DCAF5 EIF1 PTP4A1 H2AFJ NAMPT NOP56 TBPL1 NBR1 PALLD DCAF12 SEPHS1 MIER1 BIRC2 HIST2H2BE HSF2 LRRC58 |
| **Branch 5** |
| AGPAT2 CAPN2 MARCKS CDH1 CD82 TMED6 USH1C LLGL2 ACTN1 SLC9A3R1 CARHSP1 ARHGEF10L ABHD15 TMEM176A TMEM176B EPS8L2 NCOA6 PXK ADCK3 IGFBP7 KIAA2018 MLLT10 SETD2 WASL SLC4A2 VEZT GNB2 DPP8 PPHLN1 IFT46 ARAF WDR55 GADD45G UBE2J2 AAAS TRMT6 MAX PLOD1 PFKL GNL3 MAPKAPK5 APCDD1L-AS1 DHRS3 PLA2G16 TUBGCP4 PREP C9orf64 CDC73 SRPK2 TIMM22 SYT4 C9orf156 PHF23 FNDC3A ODF2L PTMS OBSL1 DMKN TXLNA USP8 ZNF346 |
| **Branch 6** |
| TPI1 PGAM1 BNIP3L ENO1 EDARADD RNF11 HMGCL TOLLIP ARHGDIA UBE2I AAMP ALDOC C14orf1 GCG C4orf3 GC TMCO1 CLNS1A PROSC MRPL49 PITHD1 POLD2 DHPS DNAJA3 DCK MRPS16 PGLS APMAP CCT7 C11orf73 DLGAP4 TALDO1 BROX CCS CCNH NDUFV1 PMVK VAMP8 PYCR2 ANXA6 SPRYD3 NHP2 ARFIP2 FTO PAICS DDX17 THTPA MRPL48 NDUFB5 LYRM9 ACADM UBE2N DARS2 |
| **Branch 7** |
| PDCD4 HMGN3 PKM MOCS2 ATP1B1 PAX6 C10orf10 TNFRSF1A SPINT1 GPR56 MAGED2 LGMN BEX4 CD47 TSPAN31 ANXA11 COPS7A AP1G2 NPTN ALDH1A1 RGS4 ARL15 CAMK2N1 TMED7 SCAMP5 ID3 ID1 ID2 SAMD11 CDKN1A ID4 SMAD7 SKIL BTG2 DBI EGR1 FOS JUNB CHRAC1 KLF10 MORN2 DUSP1 ISL1 PLK2 RHOBTB3 IER3 DDR1 LINC00673 CDC42BPA TMEM30B CASC4 TK2 WDR6 RNF220 C7orf49 MAPK1IP1L GSTO1 MICU2 MDH1 CISD3 GSTO2 NME3 GRSF1 COMT PAFAH1B3 DYNLL1 DYNLT1 RNASET2 NIT1 FAM219B PSMG3 MRPL12 ARHGEF3 TMED1 TMEM175 UXT OSTF1 BLOC1S1 C15orf57 PFDN4 B9D1 TUBGCP5 |
| **Branch 8** |
| TIMM23 PSMB3 ATP6V0E2 LGALS3BP NEUROD1 HEPACAM2 BMP5 RASD1 CASR LDOC1 ACLY MAFB TMEM37 PAPSS2 RIN2 TSC22D1 MAFA NPTX2 RASGRF1 PFKFB2 LMO1 CD14 RPH3AL PDX1 GCK RAB11B FFAR1 SLC25A34 SNX14 ABHD14A CDIPT RPN2 GPAA1 RAB1B SERPINB1 ACAT1 THYN1 NCSTN CYP27A1 RNF130 PLP2 DPM3 |

| **Branch 9** |
| --- |
| AP1M2 HCFC1R1 ECHS1 GLOD4 TIMM50 BBX CREBL2 PCYOX1 DPY30 PRMT6 RNASEH2C HIST1H4C RAP1GAP2 PDK3 NKX2-2 RAP1GAP NFYC COA5 DCTN6 NIPSNAP1 IPO9 AP1S1 SLC25A39 TUBA1B TUBA1A NDUFAF3 LDLRAP1 SLC35F6 SCOC CNOT7 CHURC1 HAT1 MRPS11 LYRM5 C5orf24 ACO2 PSMA3 CBX3 COMMD8 EEF1E1 BLVRA RAB6A DAP3 TIMMDC1 MED29 PUF60 MRPL15 HAX1 GOT2 NIPSNAP3A |
| **Branch 10** |
| NAPA TRAPPC1 CHMP5 HSBP1 SEPW1 ENO2 PGK1 TMEM14B COX5A SSBP1 NDUFB10 RAB5C LSM4 ANAPC11 WBP2 NDUFB4 TMEM141 PRDX5 NDUFB1 C14orf2 MRPL33 COMMD6 COX5B TOMM22 COX7A2 COX17 NDUFA7 ROMO1 UQCRQ COX6C PRDX2 ZCRB1 ATPIF1 PCP4 SCP2 FKBP3 C17orf89 PPIA SERPINB6 HMGN2 SRP9 POLR2I MRPL20 UROS FIS1 SUMO3 PRMT2 GTF3C6 OAZ2 NDUFB3 RBX1 S100A11 MRPL53 MPV17 OCEL1 VPS29 MGST3 PTGR1 TCEAL8 PARK7 COX14 SNRPE RAB3A SNRPD3 C18orf32 GNG4 VPS72 DLD NDUFB11 C1D ARL2 PSMB1 SUMO2 ATP5O MYL12A PSMA1 NDUFA8 TIMM17B NGRN TXN PSMA6 PSMC2 GTF2A2 HMGN1 COX7A2L CSNK1A1 RAB1A NUDCD2 ATP5H VCP MPC2 PCBD1 C11orf31 EIF2AK1 NTPCR GRHPR ERP29 CNP POLR2E TSR2 ATOX1 ARL3 POLR2L LAMTOR1 ATP5G3 ALKBH7 LINC00263 PSMB5 METTL5 CMC1 EDF1 |
| **Branch 11** |
| BCKDK PLSCR1 UBA2 IDH2 MRPS2 CCNL1 HNRNPDL H3F3C PAFAH1B1 MAPRE1 CHMP1B SYPL1 FNTA ZNF24 STX5 WDR48 C11orf58 TMEM134 TMBIM1 RQCD1 ARL4D RBP4 KCTD6 FAM3C TMEM18 RPS27L ITGB3BP EXOSC1 TRAPPC4 SENP7 TRAPPC6B TRIM52-AS1 HCG18 ZFPL1 EBNA1BP2 ANP32B PTPN3 BCL2L13 DDX50 NELFE CHCHD7 DARS PFKP UBE3A PSMB2 DDX3X MPHOSPH10 SMEK2 SMARCA5 ZBED5 EPB41 HNRNPUL1 GATAD1 VMA21 DYNC1LI1 LINC01420 TTC28-AS1 MPLKIP ZNF585A BTG1 VPS4B PDCD6IP EIF5A EIF5AL1 TTC9C POLE3 POLR2H TMEM68 SHMT2 LUC7L2 LSM7 GINM1 USP30 DERA WARS ARL1 FLOT2 |

Table S2: NDCGs in alpha cells.

| |
|---|
| **Branch 1** |
| SCG2 PAM ATP1B1 CHGB CPE GPX3 PDK4 MUC13 CLDN7 RPN2 SURF4 STK19 RGS9 TMED10 PRDX4 SLC35B1 RHBDD2 SEC24D CXCL16 NT5DC1 MAGT1 C1GALT1C1 TMED9 SAR1A PDGFRL HSP90B2P HSP90B1 PDIA4 P4HB CALR ALG11 PDIA6 CTSL TMED3 PLOD3 MYDGF EXT2 SLC39A11 SEZ6L FKBP11 ATP6AP2 SORBS2 BTF3L4 PSMD5 DGUOK IER3IP1 |
| **Branch 2** |
| ALDOA TPI1 PGK1 GAPDH ENO1 LDHA PLOD2 PFKP KL MTFP1 S100A10 MIF-AS1 NDRG1 SYNPO P4HA1 SCD SEC61G C4orf3 EDARADD ALDOC PDK1 CFC1 ENO2 TMEM45B ZNF395 PFKL PDCD4 ARRDC4 BNIP3 STC2 RSL24D1 PAFAH1B1 TSC22D1 TBRG1 |
| **Branch 3** |
| SNAP25 ASAH1 TM9SF2 PSMA2 ANXA7 SRI GHITM CALM2 CLIC1 LMCD1 CNIH1 CSDE1 UNC50 ATP6V1B2 DSTN ATP5C1 TMED2 NPTN PRDX3 YWHAQ SNX3 PPP1CB PAIP2 MYL12B ZNF706 H3F3B H3F3C BTF3 CNBP SRP9 C11orf58 VDAC1 DDX24 SARS PSMD8 VDAC2 POMP LMBRD1 GNG10 EIF3H RBM4 HDAC2 TMBIM6 H2AFZ TM9SF1 TMX2 EPCAM SARAF ITM2B COMMD8 VMP1 RAP1B ITGB1 TMX1 HSPA8 YIF1A CCT5 PTGES3 PSMA3 EIF4G2 REEP5 TMOD1 RAB2A PSMB1 YIPF6 TMF1 ANXA5 RTN3 AAMP ARF1 MRFAP1 MAPRE1 TAGLN2 NDUFA9 TCP1 TMEM50A VPS29 TMEM59 QPCT HM13 TSPAN3 C5orf15 PDIA3 PDIA3P1 RAN TMEM230 ATIC C1QBP HADHB DLD PPA1 HBS1L GOT2 RER1 NAE1 TMEM179B TMEM208 TM2D3 ADSL SLC25A4 HSPA9 SLC25A3 ERP44 UQCRFS1 MRPL13 CD46 CD164 MAP1LC3B2 MAP1LC3B GNS MGST3 VAPA CLTA RAB7A AP2M1 TMEM33 SCOC PAPOLA MRPL36 NDUFB9 MMADHC MOB4 YWHAH HSPB1 CACYBP TMEM163 SYT5 BZW1 TVP23B C14orf119 |
| **Branch 4** |
| WDR83OS BEX2 FKBP2 RPS9 H2AFJ RPS4X RPS12 RPLP2 RPL23A RPL12 UBA52 RPL34 POLR2L OST4 NDUFB3 TMEM258 COX5B TRMT112 NDUFA2 ATPIF1 NDUFA3 UQCRQ UQCR11 SHFM1 COX8A GABARAP USMG5 NDUFA1 ATP5I ROMO1 RPS11 TCEB2 SERF2 COX6C ATP5G2 UBL5 COX17 NDUFB4 RPL27 SNRPD2 SLIRP NDUFB1 ATP5L COX7B ATP5E ATP5EP2 COX7C RPL35A RPL24 RPL23 TPT1 RPL41 TMA7 ATP5J2 RPL13 RPL27A RPL37A RPL31 RPS14 RPLP1 RPS24 RPS18 RPL26 RPL11 RPS13 RPS19 RPS25 RPL30 RPL37 RPS15 RPL36 RPLP0 RPL8 EEF1A1 RPL3 RPS6 RPL14 FTH1 FTL RPS3 RPS23 RPL5 RPL19 RPL38 PEBP1 GPX4 RPL18 RPL13AP5 RPL13A GUK1 DRAP1 FIS1 ATP6V1E1 RPS7 C14orf2 |

Table S3: GO-terms in beta cell NDCGs.

| Group | GO term | p value |
|---|---|---|
| CC | CYTOSOLIC RIBOSOME | 3.1e-57 |
| BP | COTRANSLATIONAL PROTEIN TARGETING TO MEMBRANE | 1.7e-54 |
| BP | ESTABLISHMENT OF PROTEIN LOCALIZATION TO ENDOPLASMIC RETICULUM | 4.1e-52 |
| BP | NUCLEAR TRANSCRIBED MRNA CATABOLIC PROCESS NONSENSE MEDIATED DECAY | 9.5e-52 |
| BP | PROTEIN LOCALIZATION TO ENDOPLASMIC RETICULUM | 4.1e-49 |
| MF | STRUCTURAL CONSTITUENT OF RIBOSOME | 7.7e-48 |
| BP | PROTEIN TARGETING TO MEMBRANE | 1.1e-47 |
| BP | VIRAL GENE EXPRESSION | 1.7e-45 |
| CC | RIBOSOMAL SUBUNIT | 2.1e-45 |
| BP | TRANSLATIONAL INITIATION | 1.4e-44 |
| CC | RIBOSOME | 4e-44 |
| BP | NUCLEAR TRANSCRIBED MRNA CATABOLIC PROCESS | 8.2e-44 |
| CC | CYTOSOLIC LARGE RIBOSOMAL SUBUNIT | 2.8e-40 |
| BP | ESTABLISHMENT OF PROTEIN LOCALIZATION TO MEMBRANE | 8.2e-40 |
| BP | PROTEIN TARGETING | 2.9e-38 |
| MF | STRUCTURAL MOLECULE ACTIVITY | 1.1e-37 |
| BP | RNA CATABOLIC PROCESS | 2.8e-37 |
| BP | CELLULAR NITROGEN COMPOUND CATABOLIC PROCESS | 1e-33 |
| BP | ORGANIC CYCLIC COMPOUND CATABOLIC PROCESS | 4.1e-33 |
| BP | ESTABLISHMENT OF PROTEIN LOCALIZATION TO ORGANELLE | 1.5e-32 |
| BP | PROTEIN LOCALIZATION TO MEMBRANE | 2.5e-32 |
| CC | LARGE RIBOSOMAL SUBUNIT | 2.5e-30 |
| BP | PEPTIDE BIOSYNTHETIC PROCESS | 1.5e-29 |
| CC | RIBONUCLEOPROTEIN COMPLEX | 2.5e-29 |
| BP | PEPTIDE METABOLIC PROCESS | 8.4e-29 |
| BP | SYMBIOTIC PROCESS | 2.5e-28 |
| BP | PROTEIN LOCALIZATION TO ORGANELLE | 2.9e-28 |
| BP | AMIDE BIOSYNTHETIC PROCESS | 1.9e-27 |
| BP | CELLULAR AMIDE METABOLIC PROCESS | 1.6e-25 |
| BP | INTRACELLULAR PROTEIN TRANSPORT | 9.8e-25 |
| BP | MRNA METABOLIC PROCESS | 1.6e-24 |
| BP | MACROMOLECULE CATABOLIC PROCESS | 1.9e-23 |
| BP | CELLULAR MACROMOLECULE CATABOLIC PROCESS | 5.3e-23 |
| BP | ORGANONITROGEN COMPOUND BIOSYNTHETIC PROCESS | 1e-22 |
| BP | INTRACELLULAR TRANSPORT | 7.3e-22 |
| CC | POLYSOMAL RIBOSOME | 1.9e-21 |
| CC | CELL SUBSTRATE JUNCTION | 1.5e-20 |
| BP | CELLULAR MACROMOLECULE LOCALIZATION | 3.4e-20 |
| CC | POLYSOME | 7.2e-19 |
| BP | CYTOPLASMIC TRANSLATION | 1.7e-17 |
| MF | RNA BINDING | 3.3e-17 |
| CC | ANCHORING JUNCTION | 3.1e-16 |
| CC | CYTOSOLIC SMALL RIBOSOMAL SUBUNIT | 4.8e-16 |
| MF | RRNA BINDING | 1.5e-13 |
| BP | RIBOSOME BIOGENESIS | 3.6e-13 |
| CC | SMALL RIBOSOMAL SUBUNIT | 4.1e-13 |
| BP | RIBOSOMAL LARGE SUBUNIT BIOGENESIS | 2.1e-12 |
| BP | RIBOSOME ASSEMBLY | 1.2e-11 |
| BP | RIBONUCLEOPROTEIN COMPLEX BIOGENESIS | 3e-11 |
| CC | SYNAPSE | 5.2e-10 |
| BP | RRNA METABOLIC PROCESS | 1.9e-08 |
| CC | NEURON TO NEURON SYNAPSE | 8e-08 |
| BP | RIBONUCLEOPROTEIN COMPLEX SUBUNIT ORGANIZATION | 1.3e-07 |
| BP | NCRNA PROCESSING | 1.4e-06 |
| CC | POSTSYNAPSE | 2.2e-06 |
| BP | NCRNA METABOLIC PROCESS | 8e-06 |
| CC | NUCLEOLUS | 1.3e-05 |
| BP | ORGANELLE ASSEMBLY | 4.2e-05 |

**GO terms detected in branch 1.**

| Group | GO term | p value |
|---|---|---|
| BP | SECRETION | 2.2e-05 |
| BP | CELL CELL SIGNALING | 0.00019 |
| BP | HOMEOSTATIC PROCESS | 0.00098 |

**GO terms detected in branch 2.**

| Group | GO term | p value |
|---|---|---|
| CC | VACUOLE | 2.4e-08 |
| CC | VACUOLAR MEMBRANE | 7e-08 |
| CC | WHOLE MEMBRANE | 1.3e-07 |
| CC | ENDOSOME | 4.1e-05 |
| CC | ENDOPLASMIC RETICULUM | 5.9e-05 |
| BP | EXOCYTOSIS | 0.00019 |
| CC | SECRETORY VESICLE | 0.00027 |
| BP | SECRETION | 0.00029 |
| CC | NUCLEAR OUTER MEMBRANE ENDOPLASMIC RETICULUM MEMBRANE NETWORK | 0.0012 |
| CC | GOLGI APPARATUS | 0.0022 |

**GO terms detected in branch 3.**

| Group | GO term | p value |
|---|---|---|
| BP | ORGANIC ACID METABOLIC PROCESS | 0.00064 |
| BP | DEFENSE RESPONSE | 0.002 |

**GO terms detected in branch 4.**

| Group | GO term | p value |
|---|---|---|
| MF | CYTOSKELETAL PROTEIN BINDING | 1e-04 |
| BP | CYTOSKELETON ORGANIZATION | 0.0027 |

**GO terms detected in branch 5.**

| Group | GO term | p value |
|---|---|---|
| BP | SMALL MOLECULE BIOSYNTHETIC PROCESS | 7.8e-05 |
| BP | GENERATION OF PRECURSOR METABOLITES AND ENERGY | 0.00011 |
| BP | OXIDATION REDUCTION PROCESS | 0.00019 |
| BP | ORGANOPHOSPHATE METABOLIC PROCESS | 0.00052 |
| BP | CARBOHYDRATE DERIVATIVE METABOLIC PROCESS | 0.0027 |
| MF | IDENTICAL PROTEIN BINDING | 0.0041 |
| MF | RIBONUCLEOTIDE BINDING | 0.0042 |

**GO terms detected in branch 6.**

| Group | GO term | p value |
|---|---|---|
| BP | REGULATION OF CELL POPULATION PROLIFERATION | 1.7e-09 |
| BP | NEGATIVE REGULATION OF CELL POPULATION PROLIFERATION | 3.7e-08 |
| BP | CIRCULATORY SYSTEM DEVELOPMENT | 6.4e-08 |
| BP | NEGATIVE REGULATION OF MULTICELLULAR ORGANISMAL PROCESS | 6.6e-08 |
| BP | NEGATIVE REGULATION OF DEVELOPMENTAL PROCESS | 1.5e-07 |
| BP | NEGATIVE REGULATION OF CELL DIFFERENTIATION | 1.7e-07 |
| BP | NEGATIVE REGULATION OF CELL DEVELOPMENT | 5.9e-07 |
| BP | VASCULATURE DEVELOPMENT | 2e-06 |
| BP | REGULATION OF CELL DEVELOPMENT | 1.1e-05 |
| BP | POSITIVE REGULATION OF MULTICELLULAR ORGANISMAL PROCESS | 5e-05 |
| BP | POSITIVE REGULATION OF CELL POPULATION PROLIFERATION | 9.8e-05 |
| BP | TUBE DEVELOPMENT | 0.00012 |
| BP | REGULATION OF CELL DIFFERENTIATION | 0.00014 |
| BP | POSITIVE REGULATION OF DEVELOPMENTAL PROCESS | 0.00014 |
| BP | REPRODUCTION | 0.00017 |
| BP | TUBE MORPHOGENESIS | 0.00021 |
| BP | RESPONSE TO ENDOGENOUS STIMULUS | 0.00023 |
| BP | REGULATION OF NERVOUS SYSTEM DEVELOPMENT | 0.00036 |
| BP | CELL MOTILITY | 0.00036 |
| BP | EPITHELIUM DEVELOPMENT | 0.00043 |

**GO terms detected in branch 7.**

| Group | GO term | p value |
|---|---|---|
| BP | CELL CELL SIGNALING | 1.9e-05 |
| BP | SECRETION | 8.7e-05 |
| BP | CARBOHYDRATE DERIVATIVE METABOLIC PROCESS | 0.00051 |
| BP | RESPONSE TO OXYGEN CONTAINING COMPOUND | 0.00077 |
| BP | POSITIVE REGULATION OF SIGNALING | 0.0013 |
| BP | REGULATION OF TRANSPORT | 0.0017 |
| BP | HOMEOSTATIC PROCESS | 0.0024 |

**GO terms detected in branch 8.**

| Group | GO term | p value |
|---|---|---|
| CC | MITOCHONDRION | 9.7e-05 |
| CC | ENVELOPE | 0.0069 |

**GO terms detected in branch 9.**

| Group | GO term | p value |
|---|---|---|
| BP | OXIDATIVE PHOSPHORYLATION | 1.9e-15 |
| BP | ATP METABOLIC PROCESS | 1.6e-12 |
| CC | MITOCHONDRIAL ENVELOPE | 5.8e-12 |
| BP | ATP SYNTHESIS COUPLED ELECTRON TRANSPORT | 2.8e-11 |
| BP | RESPIRATORY ELECTRON TRANSPORT CHAIN | 1.4e-10 |
| BP | MITOCHONDRION ORGANIZATION | 2.4e-10 |
| CC | ORGANELLE INNER MEMBRANE | 3.4e-10 |
| CC | INNER MITOCHONDRIAL MEMBRANE PROTEIN COMPLEX | 1.1e-09 |
| BP | GENERATION OF PRECURSOR METABOLITES AND ENERGY | 1.7e-09 |
| BP | ELECTRON TRANSPORT CHAIN | 1.8e-09 |
| MF | OXIDOREDUCTASE ACTIVITY | 4.4e-09 |
| CC | MITOCHONDRION | 5.4e-09 |
| BP | CELLULAR RESPIRATION | 7.2e-09 |
| CC | ENVELOPE | 8.8e-09 |
| CC | MITOCHONDRIAL PROTEIN COMPLEX | 9.5e-09 |
| BP | OXIDATION REDUCTION PROCESS | 1.2e-08 |
| CC | RESPIRASOME | 1.2e-07 |
| BP | ENERGY DERIVATION BY OXIDATION OF ORGANIC COMPOUNDS | 1.8e-07 |
| BP | PROTON TRANSMEMBRANE TRANSPORT | 1.8e-06 |
| BP | MITOCHONDRIAL RESPIRATORY CHAIN COMPLEX ASSEMBLY | 3.1e-06 |
| BP | MITOCHONDRIAL TRANSPORT | 6e-06 |
| BP | TRANSMEMBRANE TRANSPORT | 1.2e-05 |
| CC | MEMBRANE PROTEIN COMPLEX | 1.7e-05 |
| BP | CELLULAR PROTEIN CONTAINING COMPLEX ASSEMBLY | 0.00022 |
| CC | CATALYTIC COMPLEX | 0.00029 |
| BP | PROTEIN CONTAINING COMPLEX SUBUNIT ORGANIZATION | 0.00038 |

**GO terms detected in branch 10.**

| Group | GO term | p value |
|---|---|---|
| BP | PROTEIN CONTAINING COMPLEX SUBUNIT ORGANIZATION | 2.7e-05 |
| BP | REGULATION OF MITOTIC CELL CYCLE | 0.00084 |
| BP | MITOTIC CELL CYCLE | 0.001 |

**GO terms detected in branch 11.**

Table S4: Differentially expressed genes in beta cells.

| Gene | Fold Change | Adjusted p value |
|---|---|---|
| KCNAB2 | 2.91 | 2.00e-15 |
| COL6A2 | 3.84 | 3.63e-06 |
| RAMP3 | 7.12 | 3.89e-06 |
| NDRG1 | -1.89 | 8.32e-06 |
| BNIP3 | -1.51 | 8.32e-06 |
| IGFBP3 | -4.48 | 1.14e-05 |
| MSN | 1.64 | 2.74e-05 |
| SMPDL3A | -1.35 | 3.76e-05 |
| TPI1 | -1.18 | 3.92e-05 |
| ARG2 | -1.58 | 4.23e-05 |
| HILPDA | -2.98 | 5.23e-05 |
| NTNG1 | 6.55 | 5.23e-05 |
| MIR210HG | -2.65 | 5.23e-05 |
| AMOTL1 | 3.70 | 9.53e-05 |
| ZNF395 | -1.41 | 9.53e-05 |
| GABRA2 | -4.17 | 1.08e-04 |
| SLC4A4 | 3.30 | 1.35e-04 |
| NPNT | 3.28 | 1.63e-04 |
| ERO1L | -1.33 | 1.70e-04 |
| MAST1 | 4.30 | 1.70e-04 |
| ENO2 | -1.02 | 3.56e-04 |
| ASTN1 | 4.84 | 5.21e-04 |
| P4HA1 | -1.49 | 1.21e-03 |
| NTRK2 | 3.75 | 1.22e-03 |
| UNC5C | 2.17 | 1.72e-03 |
| DKK3 | 3.06 | 1.94e-03 |
| TF | -5.21 | 2.10e-03 |
| LAMA5 | 1.30 | 3.19e-03 |
| SLC2A1 | -1.36 | 3.49e-03 |
| SYT1 | 3.28 | 3.49e-03 |
| DGKB | 1.95 | 3.49e-03 |
| LSAMP | 1.33 | 3.49e-03 |
| TIMP3 | 3.40 | 3.50e-03 |
| ADAMTSL2 | 1.42 | 3.70e-03 |
| FAM163A | 6.14 | 4.44e-03 |
| TRIM52-AS1 | -1.02 | 4.59e-03 |
| TMEM130 | 1.88 | 5.15e-03 |
| FRMD4A | 2.39 | 5.36e-03 |
| ZNF827 | 1.08 | 5.49e-03 |
| TMEM132C | 4.13 | 5.49e-03 |
| MOV10 | 1.22 | 5.61e-03 |
| SAMD5 | 1.42 | 6.47e-03 |
| SMAD9 | -1.30 | 6.80e-03 |
| NEAT1 | 1.62 | 8.16e-03 |
| CITED2 | -1.85 | 8.50e-03 |
| TDRD5 | 4.04 | 9.07e-03 |
| MYOF | 3.34 | 9.07e-03 |
| APOL4 | 2.44 | 9.88e-03 |
| MBOAT4 | 2.74 | 9.88e-03 |
| FAIM2 | 3.26 | 1.02e-02 |
| CCBE1 | 4.88 | 1.08e-02 |
| ST8SIA1 | 4.03 | 1.10e-02 |
| VDR | 2.68 | 1.10e-02 |
| DARS-AS1 | -1.64 | 1.10e-02 |
| TBX2 | 4.23 | 1.10e-02 |
| PRICKLE2 | 1.86 | 1.10e-02 |

| | | |
|---|---|---|
| TSHR | 3.72 | 1.31e-02 |
| ARRDC3 | -1.25 | 1.64e-02 |
| SIPA1 | -2.53 | 1.71e-02 |
| TMEM27 | -1.37 | 1.77e-02 |
| C2orf54 | -2.71 | 2.00e-02 |
| FN1 | 1.90 | 2.00e-02 |
| PTP4A3 | 1.78 | 2.00e-02 |
| PPFIBP2 | -1.76 | 2.15e-02 |
| GOLT1A | -1.30 | 2.15e-02 |
| CIART | -1.20 | 2.16e-02 |
| ONECUT2 | 1.61 | 2.32e-02 |
| C21orf62-AS1 | -1.41 | 2.43e-02 |
| FFAR4 | -1.76 | 2.45e-02 |
| KCNH8 | 2.36 | 2.46e-02 |
| HLA-DQB1 | 3.96 | 2.46e-02 |
| TCEAL6 | 1.27 | 2.46e-02 |
| FTCD | -2.08 | 2.48e-02 |
| DPYSL3 | 1.30 | 2.48e-02 |
| NPY | 2.01 | 2.56e-02 |
| MUC1 | 2.60 | 2.69e-02 |
| GAS6 | 1.91 | 2.69e-02 |
| ARL4D | -1.32 | 2.69e-02 |
| PCOLCE2 | 5.67 | 2.69e-02 |
| NRSN1 | 1.38 | 2.90e-02 |
| MTRNR2L1 | -3.35 | 2.93e-02 |
| P3H4 | 1.04 | 2.94e-02 |
| ANKRD37 | -1.59 | 2.94e-02 |
| CTTNBP2 | 2.21 | 2.94e-02 |
| SPON2 | 1.59 | 2.95e-02 |
| ACOX2 | 3.35 | 3.15e-02 |
| SSTR5-AS1 | 1.13 | 3.27e-02 |
| PHLDB1 | 1.56 | 3.33e-02 |
| PRSS50 | 3.02 | 3.33e-02 |
| EGFEM1P | 1.56 | 3.36e-02 |
| LINC00882 | -1.24 | 3.37e-02 |
| FGD6 | 1.65 | 3.37e-02 |
| HNRNPU-AS1 | -1.28 | 3.38e-02 |
| DNA2 | 2.11 | 3.85e-02 |
| ABHD15 | -1.09 | 3.92e-02 |
| NFIB | 2.77 | 3.97e-02 |
| SH3BP4 | 2.18 | 3.98e-02 |
| SLC7A14 | -1.58 | 4.09e-02 |
| SCN3A | 1.51 | 4.13e-02 |
| UNC5B | 1.12 | 4.25e-02 |
| FRAS1 | 1.97 | 4.26e-02 |
| ERICH3 | 3.63 | 4.36e-02 |
| CD82 | -1.08 | 4.44e-02 |
| TEP1 | 1.47 | 4.54e-02 |
| MDFIC | 1.55 | 4.63e-02 |
| CHL1 | -1.40 | 4.63e-02 |
| SEPT9 | 2.23 | 4.63e-02 |
| LOC101928222 | -1.55 | 4.65e-02 |
| CGN | 1.62 | 4.72e-02 |
| TMEM59L | 1.14 | 4.73e-02 |
| A1CF | -1.29 | 4.79e-02 |
| C1R | 2.50 | 4.79e-02 |

Table S4: Differentially expressed genes in alpha cells.

| Gene | Fold Change | Adjusted p value |
|---|---|---|
| P4HA1 | -1.36 | 3.00e-03 |
| BNIP3 | -1.05 | 3.11e-03 |
| CTXN2 | -1.76 | 3.18e-03 |
| MAST1 | 3.49 | 4.20e-03 |
| LTBP4 | 1.07 | 1.04e-02 |
| CACNG4 | -1.21 | 1.14e-02 |
| C5orf38 | -1.06 | 1.14e-02 |
| AK4 | -3.67 | 1.41e-02 |
| SLC2A1 | -1.94 | 1.69e-02 |
| DPEP1 | 1.73 | 2.43e-02 |
| URB1 | 1.03 | 2.43e-02 |
| MTRNR2L1 | -3.46 | 2.43e-02 |
| NDRG1 | -1.34 | 2.55e-02 |
| BEND5 | -1.04 | 4.24e-02 |
| FABP5 | -1.07 | 4.43e-02 |
| SLC6A17 | 1.29 | 4.71e-02 |

Table S5: GO-terms in alpha cell NDCGs.

| Group | GO term | p value |
|---|---|---|
| CC | ENDOPLASMIC RETICULUM | 1.2e-09 |
| CC | NUCLEAR OUTER MEMBRANE ENDOPLASMIC RETICULUM MEMBRANE NETWORK | 7e-09 |
| CC | GOLGI APPARATUS | 3.9e-07 |
| CC | GOLGI MEMBRANE | 3.5e-05 |
| CC | SECRETORY VESICLE | 7.5e-05 |
| CC | SECRETORY GRANULE | 7.7e-05 |
| CC | VESICLE MEMBRANE | 8.8e-05 |
| CC | MEMBRANE PROTEIN COMPLEX | 0.00044 |
| BP | CARBOHYDRATE DERIVATIVE METABOLIC PROCESS | 0.0015 |

**GO terms detected in branch 1.**

| Group | GO term | p value |
|---|---|---|
| BP | NAD METABOLIC PROCESS | 8.5e-18 |
| BP | PYRUVATE METABOLIC PROCESS | 9.3e-14 |
| BP | NUCLEOTIDE PHOSPHORYLATION | 1.6e-12 |
| BP | NUCLEOSIDE DIPHOSPHATE METABOLIC PROCESS | 4.4e-12 |
| BP | GLUCOSE METABOLIC PROCESS | 2.7e-11 |
| BP | CARBOHYDRATE CATABOLIC PROCESS | 3.5e-11 |
| BP | RIBOSE PHOSPHATE METABOLIC PROCESS | 1.2e-10 |
| BP | PURINE CONTAINING COMPOUND METABOLIC PROCESS | 1.5e-10 |
| BP | MONOSACCHARIDE METABOLIC PROCESS | 4.1e-10 |
| BP | OXIDATION REDUCTION PROCESS | 5.2e-10 |
| BP | MONOCARBOXYLIC ACID METABOLIC PROCESS | 1.4e-09 |
| BP | CARBOHYDRATE METABOLIC PROCESS | 2.3e-09 |
| BP | NUCLEOBASE CONTAINING SMALL MOLECULE METABOLIC PROCESS | 5.4e-09 |
| BP | GENERATION OF PRECURSOR METABOLITES AND ENERGY | 5.8e-09 |
| BP | ATP METABOLIC PROCESS | 1.3e-08 |
| BP | ORGANOPHOSPHATE METABOLIC PROCESS | 8.4e-08 |
| BP | ORGANIC ACID METABOLIC PROCESS | 1.5e-07 |
| BP | CARBOHYDRATE DERIVATIVE METABOLIC PROCESS | 1.3e-06 |
| MF | IDENTICAL PROTEIN BINDING | 0.00012 |

**GO terms detected in branch 2.**

| Group | GO term | p value |
|---|---|---|
| CC | WHOLE MEMBRANE | 6.5e-07 |
| CC | MEMBRANE PROTEIN COMPLEX | 9.1e-07 |
| CC | MITOCHONDRIAL PROTEIN COMPLEX | 4.2e-06 |
| MF | PROTEIN HETERODIMERIZATION ACTIVITY | 1.8e-05 |
| CC | SECRETORY VESICLE | 2.5e-05 |
| BP | OXIDATIVE PHOSPHORYLATION | 2.6e-05 |
| BP | MEMBRANE ORGANIZATION | 2.7e-05 |
| BP | INORGANIC ION TRANSMEMBRANE TRANSPORT | 4.3e-05 |
| CC | SECRETORY GRANULE | 6.6e-05 |
| BP | ION TRANSPORT | 8.8e-05 |
| CC | VACUOLAR MEMBRANE | 0.00011 |
| BP | CATION TRANSPORT | 0.00012 |
| CC | SECRETORY GRANULE MEMBRANE | 0.00014 |
| CC | INNER MITOCHONDRIAL MEMBRANE PROTEIN COMPLEX | 0.00014 |
| CC | ENVELOPE | 0.00014 |
| CC | CELL SURFACE | 0.00016 |
| CC | MITOCHONDRIAL ENVELOPE | 0.00017 |

**GO terms detected in branch 3.**

| Group | GO term | p value |
|---|---|---|
| CC | CYTOSOLIC RIBOSOME | 2.5e-61 |
| BP | COTRANSLATIONAL PROTEIN TARGETING TO MEMBRANE | 8.8e-61 |
| BP | ESTABLISHMENT OF PROTEIN LOCALIZATION TO ENDOPLASMIC RETICULUM | 8e-58 |
| BP | NUCLEAR TRANSCRIBED MRNA CATABOLIC PROCESS NONSENSE MEDIATED DECAY | 3.7e-57 |
| BP | PROTEIN LOCALIZATION TO ENDOPLASMIC RETICULUM | 1.4e-53 |
| MF | STRUCTURAL CONSTITUENT OF RIBOSOME | 2.9e-52 |
| BP | PROTEIN TARGETING TO MEMBRANE | 4.1e-52 |
| BP | VIRAL GENE EXPRESSION | 3.2e-50 |
| CC | RIBOSOMAL SUBUNIT | 3.8e-49 |
| BP | TRANSLATIONAL INITIATION | 9.3e-49 |
| BP | NUCLEAR TRANSCRIBED MRNA CATABOLIC PROCESS | 9.4e-46 |
| CC | RIBOSOME | 3.5e-45 |
| CC | CYTOSOLIC LARGE RIBOSOMAL SUBUNIT | 9.8e-45 |
| BP | ESTABLISHMENT OF PROTEIN LOCALIZATION TO MEMBRANE | 1.5e-43 |
| BP | PROTEIN TARGETING | 3.1e-40 |
| MF | STRUCTURAL MOLECULE ACTIVITY | 4.2e-38 |
| BP | RNA CATABOLIC PROCESS | 6.3e-36 |
| BP | ESTABLISHMENT OF PROTEIN LOCALIZATION TO ORGANELLE | 4.2e-34 |
| BP | PROTEIN LOCALIZATION TO MEMBRANE | 1.4e-33 |
| CC | LARGE RIBOSOMAL SUBUNIT | 1.8e-32 |
| BP | CELLULAR NITROGEN COMPOUND CATABOLIC PROCESS | 8.2e-32 |
| BP | PEPTIDE BIOSYNTHETIC PROCESS | 4.1e-31 |
| BP | ORGANIC CYCLIC COMPOUND CATABOLIC PROCESS | 4.3e-31 |
| BP | PEPTIDE METABOLIC PROCESS | 1.5e-30 |
| BP | AMIDE BIOSYNTHETIC PROCESS | 3.9e-28 |
| CC | RIBONUCLEOPROTEIN COMPLEX | 4.2e-28 |
| BP | ORGANONITROGEN COMPOUND BIOSYNTHETIC PROCESS | 7.2e-27 |
| BP | PROTEIN LOCALIZATION TO ORGANELLE | 1.6e-26 |
| BP | CELLULAR AMIDE METABOLIC PROCESS | 5.1e-26 |
| BP | SYMBIOTIC PROCESS | 2.7e-25 |
| BP | MRNA METABOLIC PROCESS | 1.1e-23 |
| BP | INTRACELLULAR PROTEIN TRANSPORT | 4.7e-22 |
| BP | INTRACELLULAR TRANSPORT | 6.7e-21 |
| BP | CELLULAR MACROMOLECULE CATABOLIC PROCESS | 5.1e-20 |
| BP | MACROMOLECULE CATABOLIC PROCESS | 1.3e-19 |
| CC | CYTOSOLIC SMALL RIBOSOMAL SUBUNIT | 3.9e-19 |
| BP | CYTOPLASMIC TRANSLATION | 1.5e-17 |
| MF | RRNA BINDING | 2e-17 |
| CC | SMALL RIBOSOMAL SUBUNIT | 7e-17 |
| CC | CELL SUBSTRATE JUNCTION | 2.2e-16 |
| BP | OXIDATIVE PHOSPHORYLATION | 2.7e-15 |
| BP | CELLULAR MACROMOLECULE LOCALIZATION | 2.8e-15 |
| MF | RNA BINDING | 4.1e-14 |
| CC | POLYSOMAL RIBOSOME | 5.7e-14 |
| CC | POLYSOME | 2.1e-13 |
| CC | INNER MITOCHONDRIAL MEMBRANE PROTEIN COMPLEX | 8.6e-13 |
| BP | ATP METABOLIC PROCESS | 2.1e-12 |
| BP | RIBOSOME BIOGENESIS | 5.7e-12 |
| CC | ANCHORING JUNCTION | 8.5e-12 |
| BP | ATP SYNTHESIS COUPLED ELECTRON TRANSPORT | 1.2e-11 |
| BP | RIBOSOME ASSEMBLY | 2.1e-11 |
| BP | RESPIRATORY ELECTRON TRANSPORT CHAIN | 7.8e-11 |
| BP | RIBONUCLEOPROTEIN COMPLEX BIOGENESIS | 1.2e-10 |
| BP | RIBOSOMAL LARGE SUBUNIT BIOGENESIS | 1.7e-10 |
| CC | ORGANELLE INNER MEMBRANE | 3.4e-10 |
| MF | PROTON TRANSMEMBRANE TRANSPORTER ACTIVITY | 8.3e-10 |
| BP | PROTON TRANSMEMBRANE TRANSPORT | 1.3e-09 |

| | | |
|---|---|---|
| CC | MITOCHONDRIAL PROTEIN COMPLEX | 3.9e-09 |
| CC | RESPIRASOME | 4.2e-09 |
| BP | ELECTRON TRANSPORT CHAIN | 5e-09 |
| BP | GENERATION OF PRECURSOR METABOLITES AND ENERGY | 5.6e-09 |
| CC | RESPIRATORY CHAIN COMPLEX | 8.5e-09 |
| BP | CELLULAR RESPIRATION | 2.2e-08 |
| BP | RRNA METABOLIC PROCESS | 2.8e-08 |
| BP | RIBONUCLEOPROTEIN COMPLEX SUBUNIT ORGANIZATION | 8e-08 |
| CC | NEURON TO NEURON SYNAPSE | 1.5e-07 |
| MF | MONOVALENT INORGANIC CATION TRANSMEMBRANE TRANSPORTER ACTIVITY | 2e-07 |
| BP | RNERGY DERIVATION BY OXIDATION OF ORGANIC COMPOUNDS | 5.1e-07 |
| CC | ENVELOPE | 1.5e-06 |
| CC | NUCLEOLUS | 2.1e-06 |
| BP | CELLULAR PROTEIN CONTAINING COMPLEX ASSEMBLY | 3.8e-06 |
| BP | NCRNA PROCESSING | 5.2e-06 |
| CC | MEMBRANE PROTEIN COMPLEX | 6e-06 |
| BP | MITOCHONDRION ORGANIZATION | 7e-06 |
| CC | MITOCHONDRION | 7.3e-06 |
| BP | MONOVALENT INORGANIC CATION TRANSPORT | 8.5e-06 |
| CC | POSTSYNAPSE | 9.2e-06 |
| CC | SYNAPSE | 1.2e-05 |
| MF | CATION TRANSMEMBRANE TRANSPORTER ACTIVITY | 3.2e-05 |
| BP | NCRNA METABOLIC PROCESS | 4e-05 |
| MF | OXIDOREDUCTASE ACTIVITY | 4.4e-05 |
| BP | CATION TRANSPORT | 0.00027 |
| BP | INORGANIC ION TRANSMEMBRANE TRANSPORT | 0.00036 |
| BP | PROTEIN CONTAINING COMPLEX SUBUNIT ORGANIZATION | 0.00042 |

**GO terms detected in branch 4.**
